# Mass Spectrometry-Based Characterization of Histone Post-Translational Modifications in Sugarcane

**DOI:** 10.1101/2025.03.28.646053

**Authors:** João Paulo da Luz Silva, Felipe Marcelo Almeida-Jesus, Julia Pinheiro Chagas Da Cunha, Monalisa Sampaio Carneiro, Juan Armando Casas-Mollano, Carlos Takeshi Hotta

## Abstract

Histones, the core components of nucleosomes, play an important role in chromatin structure and function through post-translational modifications (PTMs) that regulate gene expression without altering the DNA sequence. Sugarcane (Saccharum spp.), a highly polyploid and economically significant crop, has a complex genome with a limited understanding of histone PTMs. In this study, we employed mass spectrometry to characterise the PTMs of histones in internodes of field-grown commercial sugarcane, harvested at different times of the day, with a focus on histones H2A and H2B, which had not been previously studied in this context. Using a sorghum histone catalog as a reference, we identified 26 H1, 91 H2A, 70 H2B, 5 CENH3, 36 H3.1, 23 H3.3, 18 H3.3L, and 57 H4 loci in the sugarcane genome. We detected extensive PTMs, including acetylation and methylation, across histones H2A, H2B, H3, and H4. Canonical and variant histones showed distinct post-translational modification (PTM) patterns, highlighting their functional specialisation. This study provides a comprehensive catalog of sugarcane histone PTMs, offering insights into their roles in chromatin dynamics and gene regulation, with potential applications in crop improvement and stress adaptation.

## Introduction

Histones, the fundamental components of nucleosomes, play an important role in the structure and function of chromatin alongside DNA (Strahl & Allis, 2000). Each nucleosome is an octamer consisting of two copies of histones H2A, H2B, H3, and H4, wrapped by approximately 146 DNA base pairs. At its most basic level, chromatin exists as a 10-nm fiber, which can further condense with the involvement of the histone H1 and other proteins (Kouzarides, 2007). Canonical histones are rapidly synthesised and deposited behind replication forks during the S phase of the cell cycle, while non-canonical histones replace them at specific sites. This dynamic process alters chromatin structure and influences cellular functions, including replication, transcription, and the formation of heterochromatin (Henikoff & Smith, 2015).

Post-translational modifications (PTMs) of histones are essential for regulating nucleosome stability and recruiting proteins that drive epigenetic changes in gene expression, all without altering the underlying DNA sequence (Nunez-Vazquez *et al*., 2022; Joseph & Young, 2023; Vivek Hari Sundar *et al*., 2024). These modifications, primarily occurring on the N-terminal tails of histones, include lysine acetylation, methylation of lysine and arginine, and phosphorylation of serine, threonine, and tyrosine residues, among others (Moraes *et al*., 2015).

Sugarcane (Saccharum spp.) is a monocot of the Poaceae family (Cheavegatti-Gianotto *et al*., 2011). This commercially significant C4 plant, known for its high biomass accumulation and sucrose storage in the stem, possesses a highly polyploid and aneuploid genome due to interspecific hybridisations between *Saccharum officinarum* L., *S. spontaneum* L., and related species (Garsmeur *et al*., 2018). Given its economic and strategic importance, sugarcane is cultivated at various scales (Dal-Bianco *et al*., 2012).

Despite its significance, there is limited knowledge about histone PTMs in sugarcane. An H3K27 demethylase has been described as repressing drought-stress responses (Yu *et al*., 2024), whereas an H3K4 demethylase has been implicated in the regulation of sugarcane flowering (Chen *et al*., 2024). Previous research identified conserved PTMs in histones H3 and H4 in 6-month-old sugarcane leaves grown under controlled conditions (Moraes *et al*., 2015).

In this study, we employed mass spectrometry to characterise histone PTMs in the internodes of field-grown sugarcane. Samples were collected at various times of the day, enabling us to investigate temporal variation in histone modifications, an aspect often overlooked in controlled studies. We utilised a sorghum histone catalog to identify sugarcane histone genes in the recently sequenced genome and to characterise PTMs in histones H2A and H2B, which had not been previously characterised.

## Materials & Methods

### Plant Materials

Sugarcane cultivar SP80-3280 (Saccharum spp. hybrid) was grown under field conditions at the Federal University of São Carlos, campus Araras, in São Paulo state, Brazil (22°21′25″ S, 47°23′3″ W; altitude of 611 m). The soil was classified as a Typic Eutroferric Red Latosol. Harvesting took place on September 4 and 5, 2017. The internodes 5 and 6, organs where sucrose is still accumulating, were harvested from nine-month old plants. Dawn was at 6:16, and dusk was at 18:01 local time. Plants were sampled at five different time points after sunrise: ZT4 (4 h after sunrise), ZT7, ZT12, ZT17, and ZT22. At each time point, three or four pools of 10 plants, randomly selected within the field, were processed, frozen in liquid nitrogen, transported on dry ice, and stored at −80°C. In total, 19 independent samples were quantified.

### Identification of histone genes in sugarcane

We utilised a sorghum histone catalog to identify histone genes in sugarcane (Hu et al., 2022) and obtained histone sequences from the Sorghum bicolor genome version 5.1 (McCormick *et al*., 2018). We used reciprocal BLAST with the CDS sequences from sugarcane hybrid R570 genome (Healey *et al*., 2024) and with the SUCEST (Sugarcane EST Database) (Vettore *et al*., 2003). We also performed reciprocal BLAST using the sugarcane R570 genes to confirm the identified SAS (sugarcane assembled sequences) in SUCEST. The identified histones were annotated based on molecular function and homology, with specific annotations for histones H2A, H3, H4, and variants of H2B.

### Characterisation of histones post-translational modifications

Post-translational modifications (PTMs) in the samples were identified by first extracting nuclei from 40 g of sugarcane internodes using the methodology established by Bowler et al. and modified by Moraes et al. (Bowler *et al*., 2004; Moraes *et al*., 2015). Briefly, the frozen material was blended with a cold buffer (1 M hexylene glycol, 10 mM PIPES/KOH pH 7.0, 10 mM MgCl2, 10 mM 2-mercaptoethanol, 0.5 mM PMSF, 10 mM NaF, 20 mM sodium butyrate). Then, Triton X-100 was added to 0.5% concentration, the material was filtered through two layers of Miracloth, and the solution was incubated on ice for 10 min. Next, the nuclei were centrifuged for 10 min at 300 g and 4°C, the pellet was resuspended in Nuclei wash buffer (0.5 M hexylene glycol, 10 mM PIPES/KOH pH 7.0, 10 mM MgCl2, 0.2% Triton X-100, 10 mM 2-mercaptoethanol, 0.5 mM PMSF, 10 mM NaF, 20 mM sodium butyrate), centrifuged again twice more, and stored at −80°C. When all samples were prepared, the pellets were resuspended in 3 mL of a solution composed of 40% guanidine hydrochloride in phosphate buffer (pH 6.8), with 10 mM 2-mercaptoethanol and 0.1 mM PMSF. Then, 2.5 N sulfuric acid was added to a final concentration of 0.25 N, and the solution was incubated for 30 min on ice and then centrifuged at 10,000 g for 10 min at 4 °C. The supernatant was collected, and the pH was neutralised with potassium hydroxide. The cation exchange resin Biorex-70 (Bio-Rad) was used to extract bulk histones as described by Waterborg et al. (1995), which included several guanidine hydrochloride washes. Finally, samples were dialysed and concentrated using a centrifugation filter with a 3,000 Da cut-off (Amicon Merck).

Following SDS-PAGE analysis, a 20 µg aliquot of enriched histones was subjected to chemical derivatisation with propionic anhydride due to the presence of lysines at the N-terminal end (Garcia *et al*., 2007). Trypsin digestion was followed by a second propionylation step to enhance peptide hydrophobicity, thereby improving chromatographic retention. Peptide separation was performed using the NanoAcquity system (Waters Corp) and an Orbitrap Fusion Lumos mass spectrometer (Thermo Scientific). Data were acquired in a data-dependent manner, and the resulting raw data were analysed using MaxQuant software (Tyanova *et al*., 2016), with specific search parameters set to identify sugarcane histones and their PTMs. The peptide quantification was conducted using Skyline software (MacLean *et al*., 2010), and the results were statistically analysed to determine the significance of the observed modifications.

### Data-Independent Acquisition and Data Analysis

Peptide separation was performed using a NanoAcquity UPLC system (Waters Corp) with a linear gradient of solvent A (0.1% formic acid in water) and solvent B (acetonitrile with 0.1% formic acid). Each sample was initially loaded onto a nanoAcquity UPLC 2G-V/MTrap 5 µm Symmetry C18 180 µm X 20 mm trap column (Waters Corp) for 3 min at a flow rate of 7 µL/min. Following this step, peptides were separated on a nanoAcquity UPLC® BEH130 C18 1.7 µm, 100 µm x 100 mm analytical column (Waters Corp) with a flow rate of 0.4 µL/min.

Mass spectrometry data were acquired using a MAXIS 3G mass spectrometer (Bruker) equipped with a CaptiveSpray ion source (Bruker) operating at 2000 V and a gas flow rate of 3 L/min at 150°C. High-resolution MS1 scans were conducted at a resolution of 60,000, covering a mass range of 20 to 2200 m/z, followed by data-dependent MS2 scans. The MS2 spectra were acquired using collision-induced dissociation (CID).

Data-independent acquisition (DIA) data analysis was conducted using Skyline software, a widely used tool for targeted proteomics and quantitative mass spectrometry data analysis (MacLean *et al*., 2010). To facilitate peptide identification, a spectral library was imported into Skyline. This library contained pre-identified peptide spectra to match and confirm peptide sequences in the DIA data. Next, a FASTA file containing the target protein sequences of histones was loaded into Skyline. This file provided the necessary sequence information for the sugarcane histones identified on SUCEST. Digestion parameters were then configured for the enzyme trypsin, allowing for four missed cleavages, which account for incomplete digestion caused by chemical derivatisation with propionic anhydride. Skyline generated a list of target peptides based on the provided FASTA file and digestion parameters. This list was refined by excluding irrelevant sequences and peptides without precursors. Transition settings were then defined to monitor specific ion types, including b-ions, y-ions, and precursors. Charge states were set to +2 and +3 for each peptide. The DIA raw data files, obtained from the mass spectrometer, were imported into Skyline. The software automatically integrated the chromatographic peaks corresponding to each transition. The automated peak integration was reviewed, and manual corrections were made when misaligned peaks were detected. Following peak integration, the data were normalised to account for variations in sample preparation, instrument performance, or other experimental factors. Then, statistical analysis was performed to quantify peptide abundances.

Finally, the results were exported and visualised using Skyline’s built-in reporting features. Skyline’s visualisation capabilities enable a comprehensive assessment of the histone PTMs identified in the samples.

## Results

We have generated a catalog of sugarcane histone genes using a histone catalog from sorghum as a reference (Supplement Tables 1-8) (Hu *et al*., 2022). Histone gene loci were identified by reciprocal BLAST using the sequences from the *Sorghum bicolor* genome (v5.1) (McCormick *et al*., 2018) as queries to search and the commercial sugarcane hybrid R570 genome (Healey *et al*., 2024). We identified 26 H1, 91 H2A, 70 H2B, 5 CENH3, 36 H31, 23 H33, 18 H33L, and 57 H4 histone loci (Table 1). On average, 5.4 ± 0.6 (n = 8) sugarcane loci were identified for each histone locus in sorghum. The chromosomal distribution of the histones genes identified showed, in general, to be similar to their counterparts in sorghum (Supplemental Table 9). We also identified 124 histones in the SUCEST (Sugarcane EST database) (Table 1) through reciprocal BLAST using sequences from the *Sorghum bicolor* genome as queries. SUCEST was obtained from the commercial sugarcane SP80-3280 (Vettore *et al*., 2003), the same used in this experiment. Thus, these sequences were used as reference.

**Table 1.** Histone genes in sugarcane. – sugarcane genes in the Sugarcane Genome R570 and the SUCEST database were identified using a Sorghum Histone catalog as a reference. Sorghum sequences were taken from the Sorghum genome v5.1.The proportion of histone genes found in the Sorghum genome was compared with the Arabidopsis genome (Sorghum:Arabidopsis), the proportion of genes found in the Sugarcane R570 genome was compared with the Sorghum genome (Sugarcane:Sorghum), and the proportion of genes found in the SUCEST database was compared with the Sugarcane R570 genome (SUCEST:Sugarcane)

|  | H1 | H2A | H2B | CENH<br>3 | H31 | H33 | H33L | H4 |
| --- | --- | --- | --- | --- | --- | --- | --- | --- |
| **Arabidopsis** | 3 | 13 | 11 | 1 | 5 | 3 | 5 | 8 |
| **Sorghum** | 4 | 18 | 13 | 1 | 8 | 4 | 3 | 11 |
| **Sorghum:Arabido**<br>**psis** | 1.3 | 1.4 | 1.2 | 1.0 | 1.6 | 1.3 | 0.6 | 1.4 |
| **Sugarcane** | 26 | 91 | 70 | 5 | 36 | 23 | 18 | 57 |
| **Sugarcane:Sorgh**<br>**um** | 6.5 | 5.1 | 5.4 | 5.0 | 4.5 | 5.8 | 6.0 | 5.2 |
| **SUCEST** | 19 | 23 | 20 | 1 | 20 | 16 | 0 | 23 |
| **SUCEST:Sugarca**<br>**ne** | 0.7 | 0.3 | 0.3 | 0.2 | 0.6 | 0.7 | 0.0 | 0.4 |

The internodes 5 and 6 of 9-month-old commercial sugarcane (SP80 3280) grown in the field were harvested at five different times of the day to characterise sugarcane histone post-translational modifications (PTMs) using a mass spectrometer. The histone sequences in the SUCEST database were used to identify the peptides. The most abundant peptide was K9prSTGGK14acAPR, which is part of the H3 histones (Table 2). Methylation (mono-, di-, and tri-) and acetylation were the most common PTMs. Canonical and histone variants were identified, providing valuable insights into their diversity and functional specialisation in sugarcane. Specifically, we have identified and cataloged PTMs in histones H2A, H2B, H3.1, H3.3, and H4 (Table 3).

**Table 2.**
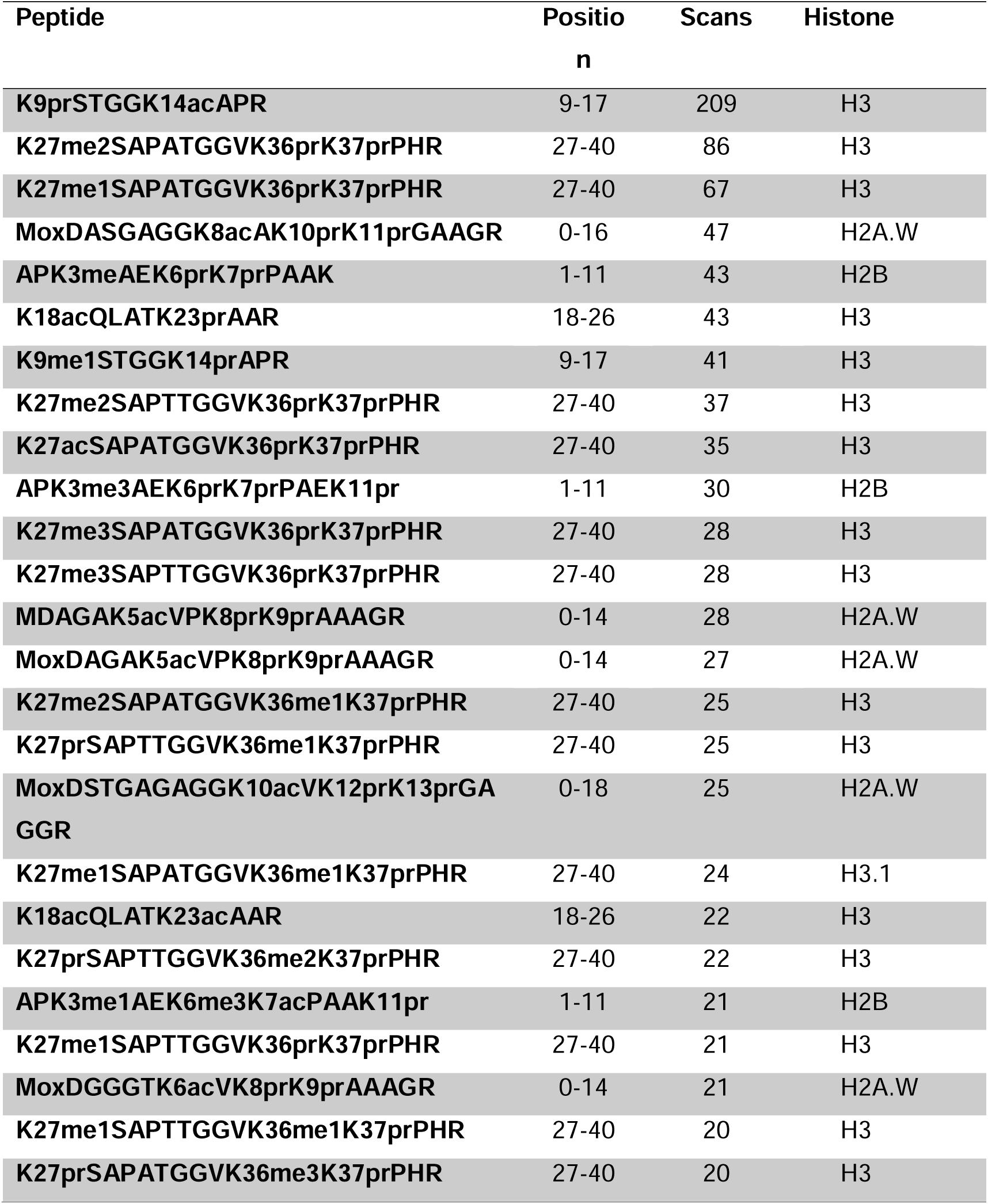
Most abundant modified peptides in sugarcane. Peptides with a scan count percentile above 0.9 were sorted in descending order based on the number of scans. The peptide modifications are represented as follows: me1 (methylation); me2 (demethylation); me3 (trimethylation); ac (acetylation); pr (propionylation); ox (oxidation).

**Table 3.**
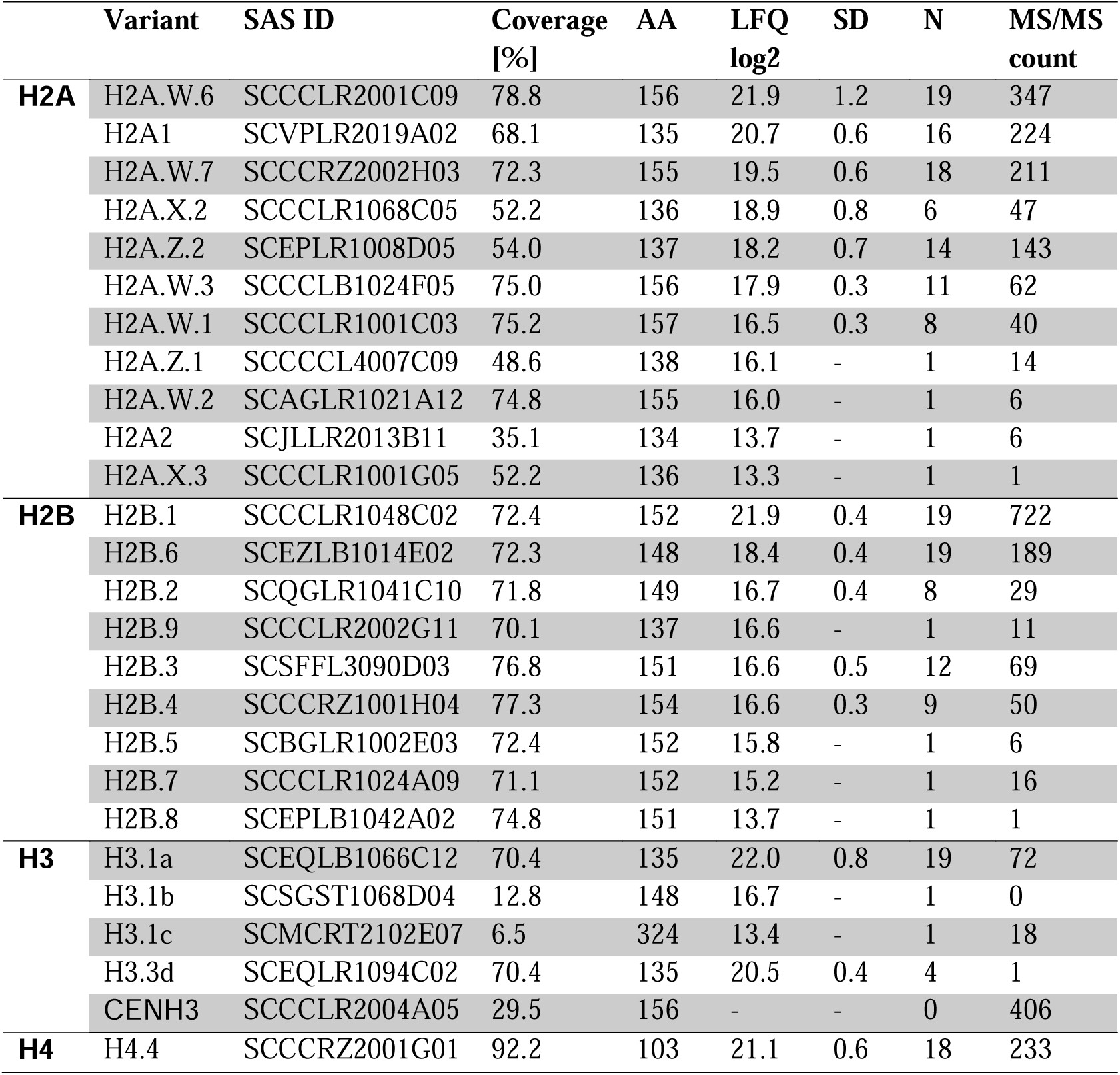
Annotation of the histones identified for internode 3-5. The acronym SAS refers to the gene deposited in the SUCEST database. “Variant” refers to the annotation for each type of histone identified. “Coverage [%]” shows the percentage of the protein’s extent identified in this study. “AA” indicates the amino acid sequence length for each histone. “LFQ log2” shows the average protein quantity, “SD” represents the standard deviation, while “N” presents the number of biological replicates for the protein quantity data. The last column shows the number of MS/MS identifications.

We have identified 27 histones and variants: 12 H2A, 9 H2B, 5 H3, and 1 H4. Approximately 63% of the identified histones had coverage greater than 70%. Only ten histones were detected in more than 50% of the 19 independent samples: 5 H2A, 3 H2B, 1 H3, and 1 H4 (Table 2). As a comparison, the SUCEST database contains 20 different H2A proteins, of which 60% were identified by mass spectrometry, 16 H2B (56%), 5 H3 (100%), and 5 H4 (20%).

We identified numerous PTM sites on histones H2A (Figures 1 and 2) and H2B (Figure 3). Canonical H2A, as well as H2A.Z and H2A.X variants, showed extensive modifications. H2A1 has four lysines (5, 13, 14, and 21) that are acetylated, mono-, di-, and tri-methylated, as well as arginine that is monomethylated (Figure 1a). H2A2 has only three lysines (5, 13, and 14), with only monomethylation and acetylation. It also contains arginine (18) with mono- or demethylation (Figure 1b). The histone variant H2A.X .2 has one lysine with trimethylation and two lysines (20 and 35) monomethylation (Figure 1c). In contrast, H2A.X .3 has three lysines with mono, trimethylation, and acetylation and one arginine with monomethylation (Figure 1d).

**Fig. 1.**
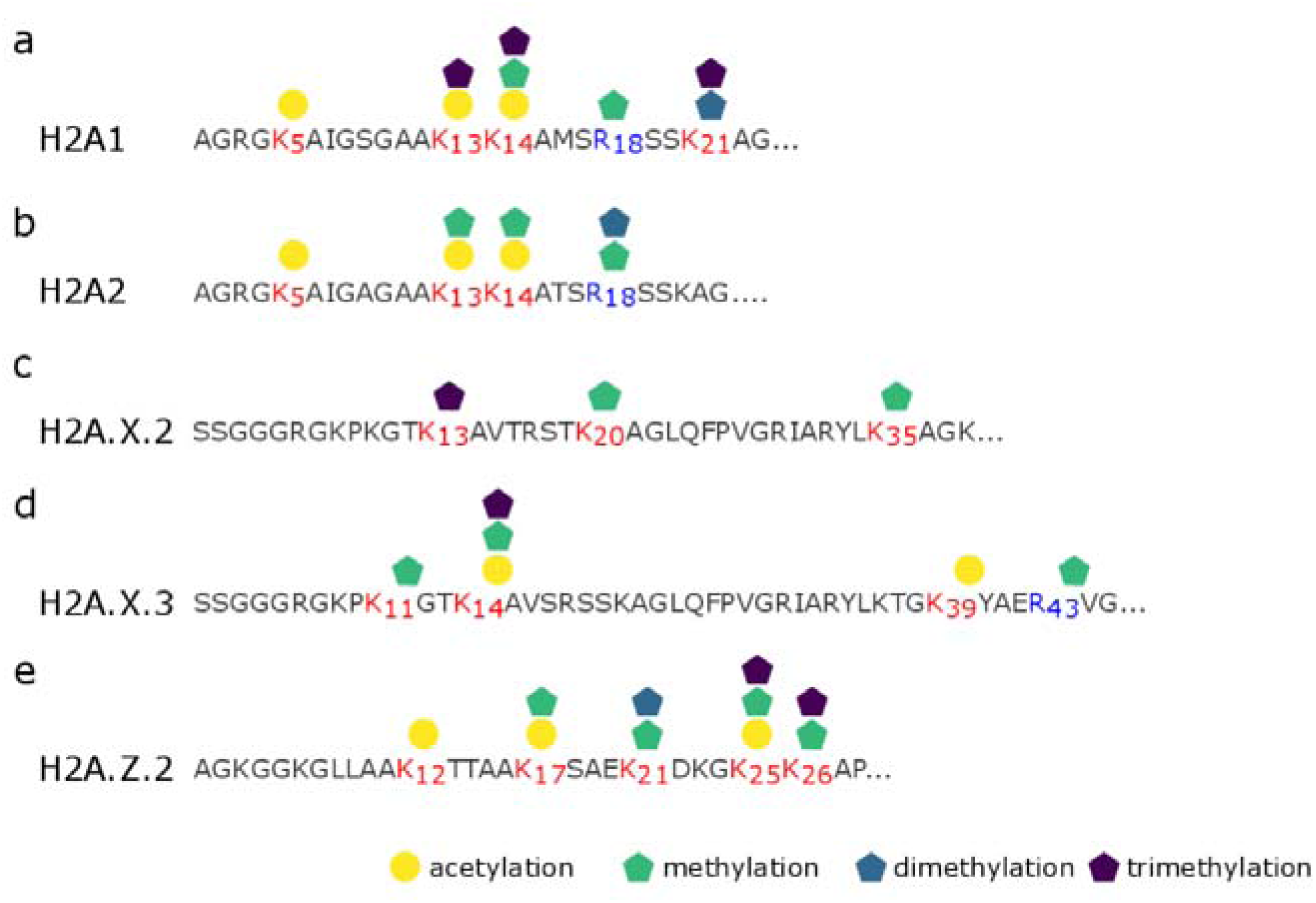
Post-Translational Modifications Identified in H2A histones variants. Illustration showing PTMs identified in **(a)** canonical H2A1, **(b)** canonical H2A2, **(c)** H2A.X.2, **(d)** H2A.X.3, and **(e)** H2A.Z.2. Amino acids are represented by their single-letter codes. Modified lysines are highlighted in red, and modified arginines are highlighted in blue, with their positions indicated by a subscripted number. Methylation (green pentagon), di-methylation (blue pentagon), tri-methylation (purple pentagon), and acetylation (yellow circle) of amino acids are indicated.

**Fig. 2.**
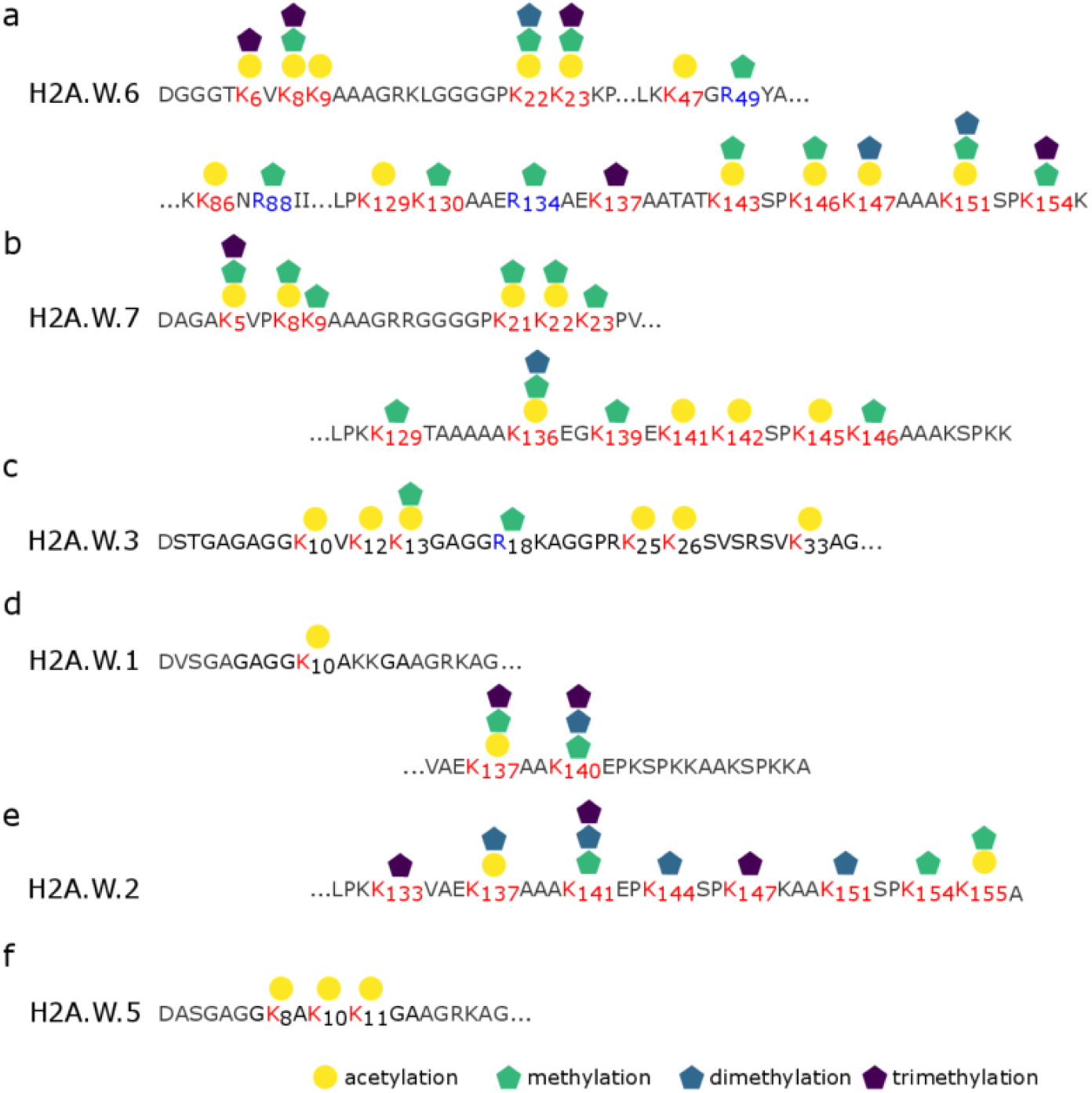
Post-Translational Modifications Identified in the histone variant H2A.W. Illustration showing PTMs identified in **(a)** H2A.W.6, **(b)** H2A.W.7, **(c)** H2A.W.3, **d)** H2A.W.1, **(e)** H2A.W.2, and **(f)** H2A.W.5. Amino acids are represented by their single-letter codes. Modified lysines are highlighted in red, and modified arginines are highlighted in blue, with their positions indicated by a subscripted number. Methylation (green pentagon), di-methylation (blue pentagon), tri-methylation (purple pentagon), and acetylation (yellow circle) of amino acids are indicated.

**Fig. 3.**
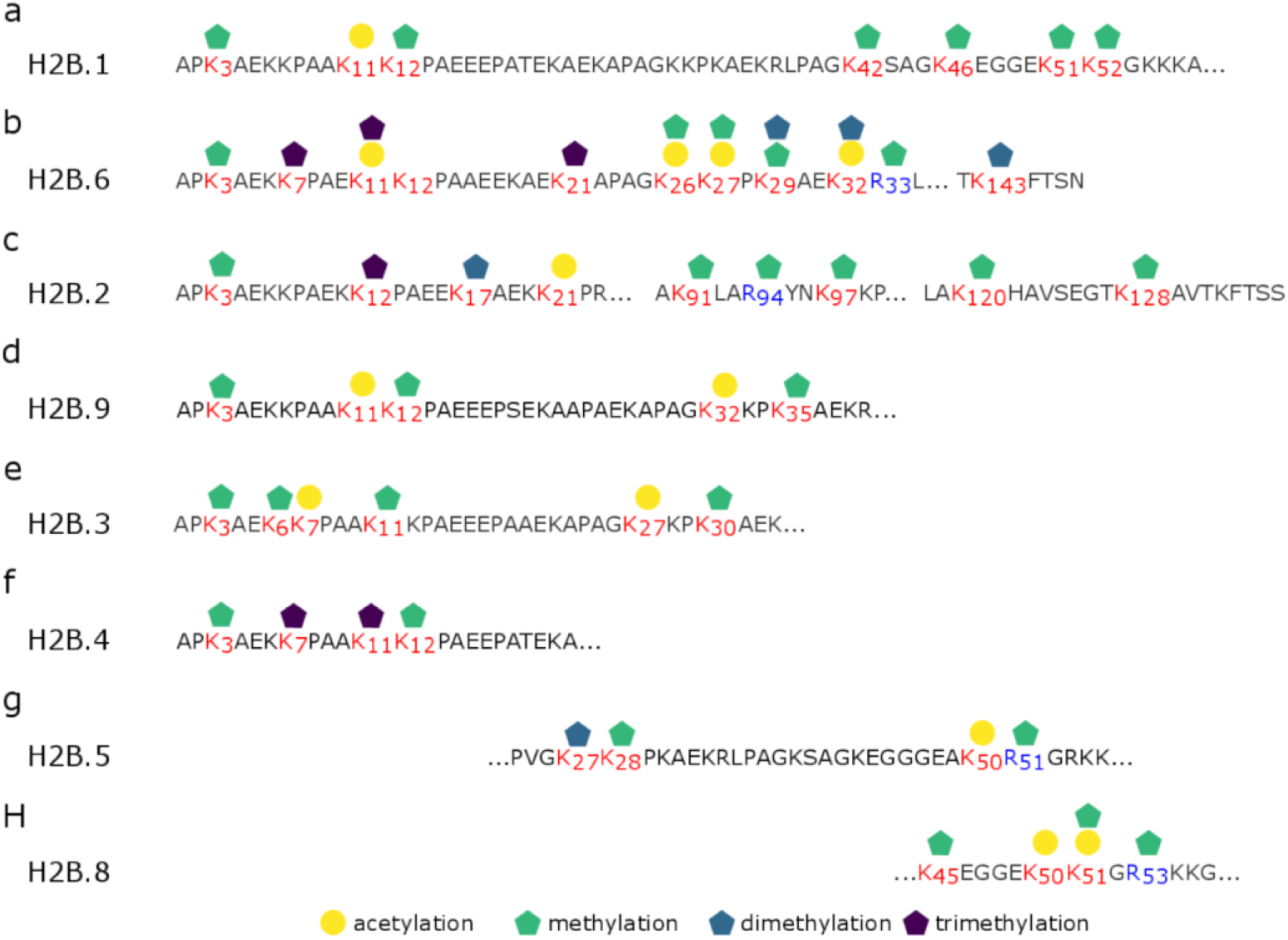
Post-Translational Modifications Identified in the histone variant H2B. Illustration showing PTMs identified in **(a)** H2B.4, **(Bb)** H2 B.6, **(c)** H2B.9, **(d)** H2B.11, **(e)** H2B.10a, **(f)** H2B.2, **(g)** H2B.10b, **(h)** H2B.10c. Amino acids are represented by their single-letter codes. Modified lysines are highlighted in red, and modified arginines are highlighted in blue, with their positions indicated by a subscripted number. Methylation (green pentagon), di-methylation (blue pentagon), tri-methylation (purple pentagon), and acetylation (yellow circle) of amino acids are indicated.

Finally, H2A.Z.2 has fine lysines with mono-, di-, and tri-methylation (Figure 1e). The histone variants H2A.W (H2A.W.6 and H2A.W.7) exhibit multiple PTMs across their N-terminal tail, central domain, and C-terminal tail, including the KSPKK domain.

H2A.W.6 has 15 lysines and three arginines with PTMs, while H2A.W.7 has 13 lysines with PTMs (Figure 2). Interestingly, in H2A, some peptides were found exclusively in their modified forms, indicating extensive modification at specific positions in sugarcane (Table 4). The N-terminal tail of H2A was particularly marked by acetylation, with H2AK5ac being the most intense PTM identified.

**Table 4.**
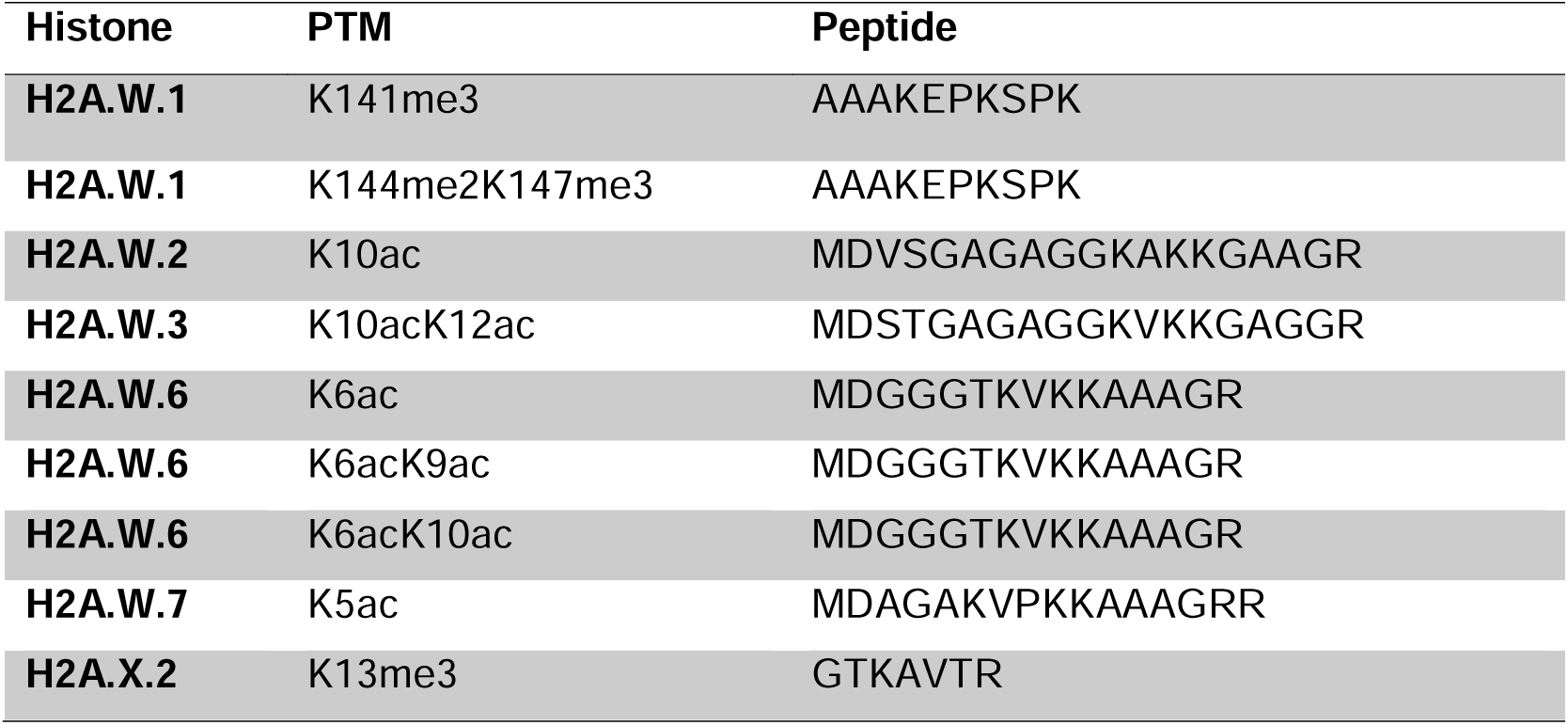
Peptides in H2A identified exclusively in their modified forms. These findings indicate that H2A is extensively modified at specific positions in sugarcane. Additionally, the N-terminal tail is prominently marked by acetylations, with H2AK5ac being the most intense PTM identified.

Histone H2B had high-frequency methylation on lysine 3, 6, 7, 11, and 12 on the N-terminal tail (Figure 3 and Table 2). PTMs were also identified between positions 21 and 52 (lysines 21, 26, 27, 29, 32, 35, 42, 46, 51, 52 and arginine 33) and in the C-terminal end (lysines 91, 97, 120, 128,143, and arginine 94) but at lower frequencies (Figure 3).

Among the previously identified histones (Moraes *et al*., 2015), histone H4 has eight lysines (8, 12, 16, 20, 31, 44, 79, and 91) with acetylation and monomethylation, and six arginines with monomethylation (17, 23, 35, 36, 78, and 92) (Figure 4). Finally, histone H3.1a has ten lysines (4, 9, 14, 18, 23, 27, 36, 37, 56, and 122) that are acetylated and subject to mono-, di-, and tri-methylation (Figure 5a). All four modifications can be found in lysine 9, 27, 36, and 37. Histone H3.1 has three arginines (at positions 8, 17, and 26) that can be mono- or dimethylated (Figure 5a). Another H3.1 variant (H3.1c) features two lysines (195 and 197) with monomethylation and four arginines (8, 14, 17, and 194) with either mono- or demethylation (Figure 5b). Histone H3.3d has three lysines (27, 36, and 37) that are acetylated and mono-, di-, and tri-methylated, as well as one arginine (40) that is mono- or dimethylated (Figure 5C). An H3L histone variant has three lysines (9, 15, and 16) that can be mono- or dimethylated (Figure 5d).

**Fig. 4.**
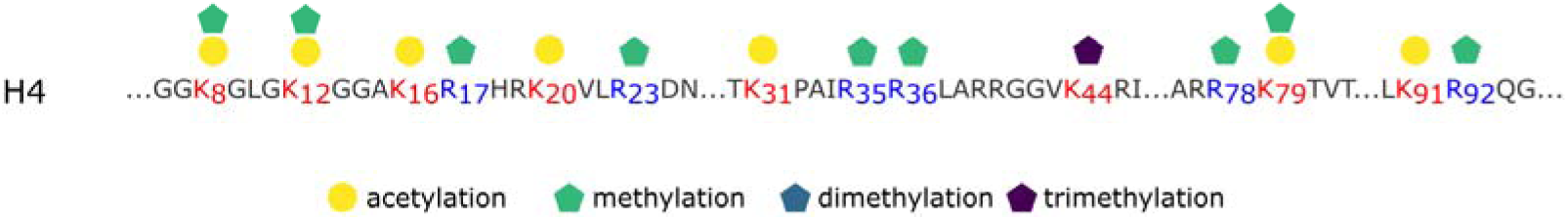
Post-Translational Modifications Identified in histones H4. Illustration of the modifications present in H4 histones. Amino acids are represented by their single-letter codes. Modified lysines are highlighted in red, and modified arginines are highlighted in blue, with their positions indicated by a subscripted number. Methylation (green pentagon), di-methylation (blue pentagon), tri-methylation (purple pentagon), and acetylation (yellow circle) of amino acids are indicated.

**Fig. 5.**
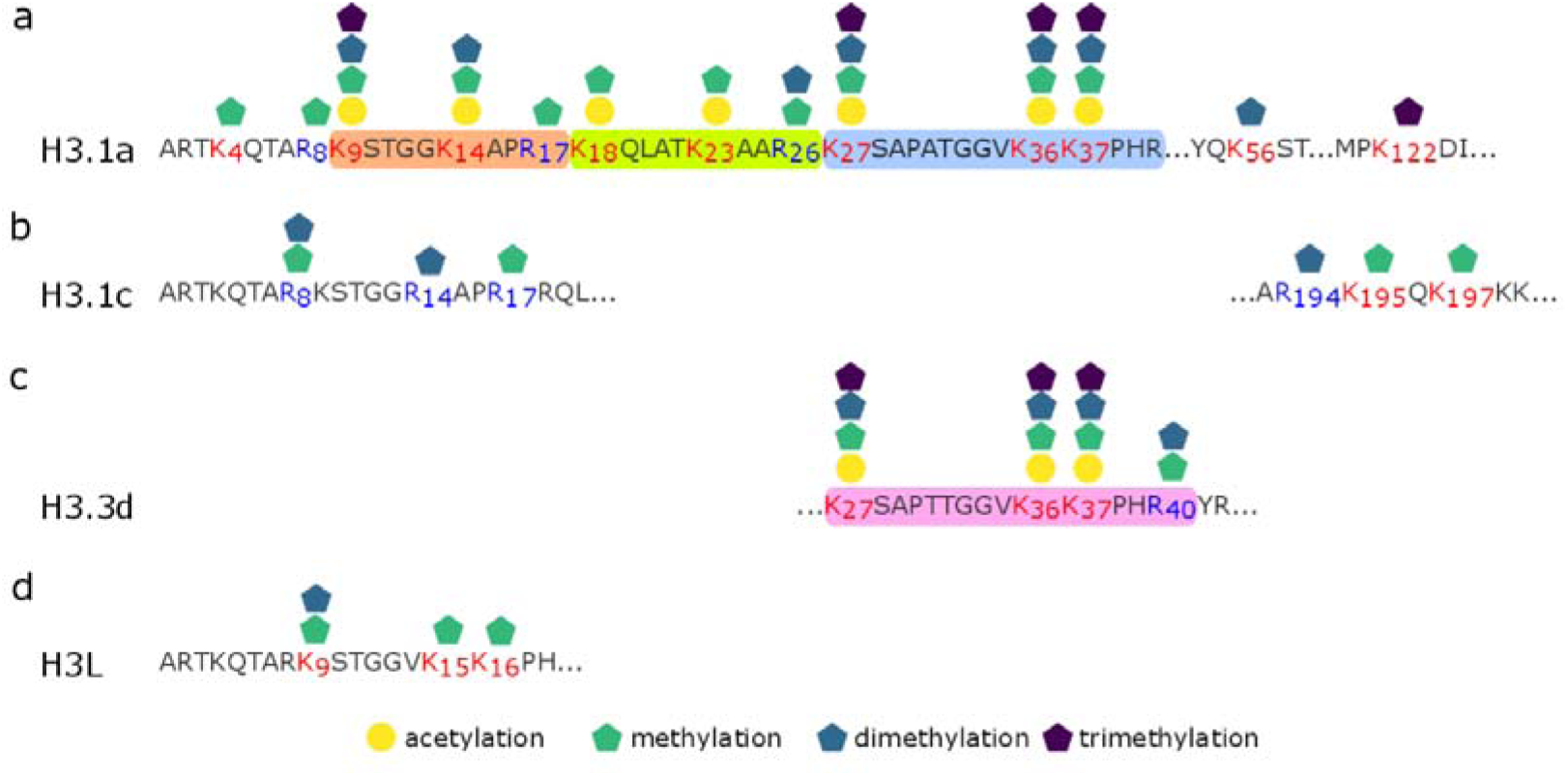
Post-Translational Modifications Identified in histones H3. Illustration of the modifications in **(a)** H3.1a, **(b)** H3.1c, **(c)** H3.3d, and **(d)** H3L histones. Amino acids are represented by their single-letter codes. Modified lysines are highlighted in red, and modified arginines are highlighted in blue, with their positions indicated by a subscripted number. Methylation (green pentagon), di-methylation (blue pentagon), tri-methylation (purple pentagon), and acetylation (yellow circle) of amino acids are indicated. Peptides 9-KSTGGKAPR-17 (orange), 18-KQLATKAAR-26 (green), 27-KSAPATGGVKKPHR-40 (blue), and 27-KSAPTTGGVKKPHR-40 (pink) were quantified using mass spectroscopy.

We could quantify specific peptides using Data-Independent Acquisition (DIA) mass spectrometry, allowing us to measure the relative abundance of modified peptides. For instance, acetylation at lysine 14 and methylation at lysine 9 were the predominant modifications in the H3.1 peptide 9-KSTGGKAPR-17, with H3.1K14ac being the most abundant form (Figure 6a). Another significant H3.1 peptide, 18-KQLATKAAR-26, exhibited high intensity in the chromatogram, with over 40% of the signal in the unmodified form (Figure 6b). Peptides 27-KSAPATGGVKKPHR-40 (H3.1) and 27-KSAPTTGGVKKPHR-40 (H3.3) exhibited distinct PMT profiles, despite being similar (Figure 6c and d).

**Fig. 6.**
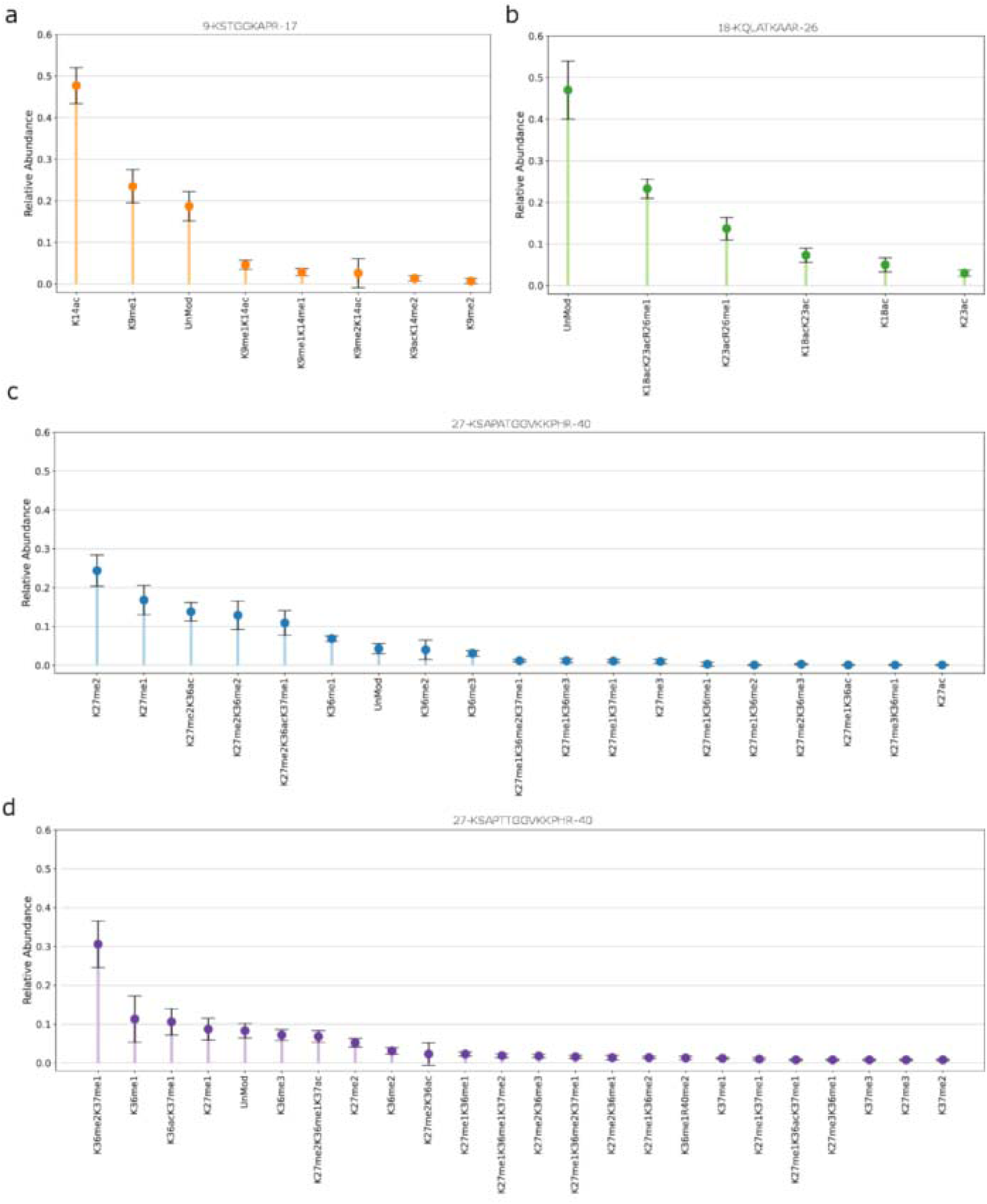
Relative abundance of post-translational modifications for histone H3 peptides. Peptides **(a)** 9-KSTGGKAPR-17, **(b)** 18-KQLATKAAR-26, **(Cc)** 27-KSAPATGGVKKPHR-40, and **(d)** 27-KSAPTTGGVKKPHR-40 were quantified using mass spectrometry. The Y-axis represents relative abundance on a scale from 0 to 1, where 0 indicates no quantity, and 1 means 100%. The bars represent the average relative abundance values, with error bars indicating the standard deviation. Each bar is labeled with its respective PTM on the X-axis, and the bar for the unmodified peptide is labeled as ‘UnMod.’

## Discussion

Histones are the main structural components of chromatin, acting dynamically to ensure regulated access to genetic information. In histones, one of the key molecular events that constantly alters chromatin organisation is the occurrence of post-translational modifications (PTMs). We identified PTMs in histones from sugarcane leaves and internodes harvested at different times of the day in the field. We employed a combination of mass spectrometry-based techniques to map the various types of modifications and their respective positions. This increased the number of known sugarcane PTMs from previous studies that only focused on histones H3 and H4 (Moraes *et al*., 2015). Most of the modifications in H3 and H4, including acetylation and methylation, were found at several sites previously described in sugarcane and Arabidopsis (Earley *et al*., 2007; Zhang *et al*., 2007). Additionally, novel modified sites were identified. Notably, this study mapped modifications in H2A and H2B for the first time in sugarcane.

Sugarcane PTMs were identified using translated SAS (sugarcane assembled sequences) from the SUCEST database (Vettore *et al*., 2003). To correctly identify the SAS, we created a catalog of sugarcane histones associating sorghum histones to sugarcane orthologs using the sugarcane R570 genome. For each sorghum gene, 5.4 histone genes were identified in the sugarcane R570 genome, which is lower than the average of 6.8 estimated using all sorghum genes (Healey *et al*., 2024). In general, the distribution of histone genes on the sugarcane chromosomes followed the same distribution found in the sorghum genome (Supplemental Table 1).

Among eukaryotes, the four core histones are highly conserved, but histone H2A has undergone significant diversification, particularly in flowering plants, where remarkable expansions of H2A variants have been observed (Kawashima *et al*., 2015). We identified 11 H2A variants belonging to the H2A.W, H2A.Z, H2A.X, and canonical H2A types in sugarcane internodes (Table 3). In different H2A variants, we found that the N-terminal tail is predominantly acetylated at lysines 5, 6, 8, 9, 13, and 14, corresponding to H2AK5ac and H2AK9ac in humans (Figures 1 and 2, Table 4). Acetylation of the N-terminal tail is described as a conserved feature across various organisms, including yeast (Xiong & Wang, 2011), humans (Tafrova & Tafrov, 2014), and Arabidopsis (Zhang *et al*., 2007). These findings suggest that this feature is also conserved in sugarcane. Histone acetylation has been previously associated with sucrose accumulation in sugarcane leaves, with acetylation detected in histone H2A, as well as in histone acetyltransferases and deacetylases, suggesting a regulatory role in sugar metabolism (Wang *et al*., 2025).

Five of the 11 histone H2A variants identified in internodes were classified as belonging to the H2A.W variant class (Figure 2 and Table 2). The H2A.W variant is characterised by the C-terminal KSPKK motif and appears exclusive to flowering plants (Kawashima *et al*., 2015). H2A.W is essential for chromatin condensation and plays a critical role in defining heterochromatin in Arabidopsis (Yelagandula *et al*., 2014). This study identified this variant as being expressed in quantities similar to, or even higher than, canonical H2A (H2A1 and H2A2) (Table 3). Considering that sugarcane has a sizeable polyploid genome of approximately 10 Gb (D’Hont *et al*., 1996; Piperidis & D’Hont, 2020), the abundance of H2A.W in both protein levels and the number of variants may be linked to the importance of heterochromatin stability. In addition to these modifications, we identified other PTMs in the C-terminal tail of H2A.W that appear less abundant (Figure 2). PTMs in the C-terminal region, particularly those involving the KSPKK motif, likely play a significant role in the function of H2A.W. This region is partially responsible for stabilising the nucleosome and interacting with histone H1, leading to increased difficulty in accessing compacted DNA (Osakabe *et al*., 2018).

H2A.X is another histone variant identified in this study. Two H2A.X histones are expressed in internodes (Table 3). H2A.X variants contain a conserved C-terminal SQEF motif in which phosphorylation of the serine occurs at DNA damage sites. H2A.X is critical in orchestrating the DNA repair pathway (van Attikum & Gasser, 2009). This study found that the PTM K13me3 is predominantly present in this histone in sugarcane (Figure 1). H2AK13 was recently identified as a site for ubiquitination, playing a role in the response to double-strand DNA breaks (Horn *et al*., 2019).

However, little is known about the methylation of this residue. Additionally, we identified the PTMs K11me3, K13me3, K14ac, K14me1, K14me3, K20me, K35me1, K39ac, and R43me1 in H2A.X (Figure 1). Given the function of this variant, these modifications may participate in DNA repair in sugarcane.

Among H2A variants in plants, H2A.Z is frequently highlighted due to its role in regulating plant responses to the environment (Kumar, 2018). However, its mechanism of action remains unclear. In *Arabidopsis*, H2A.Z exhibits opposing effects on transcription depending on its location within the gene (Sura *et al*., 2017). It has been proposed that PTMs in H2A.Z determine the role of this variant in regulating gene expression, with the amount of H2A.Z in genes being proportional to their transcriptional activity (Gómez-Zambrano *et al*., 2019). In this study, we identified two H2A.Z histones (Table 3). We also detected acetylation of lysines 12, 17, and 25, as well as varying methylation levels on lysines 17, 21, 22, and 26 (Figure 1). Acetylation of H2A.Z in the N-terminal tail is associated with transcriptional activation (Berr *et al*., 2011; Giaimo *et al*., 2019). In contrast, methylation of the N-terminal tail of H2A.Z remains poorly understood.

In addition to H2A, histone H2B was also analysed (Figure 3). H2B is classically described as the least conserved histone within the nucleosome octamer (Chabouté *et al*., 1993). In sugarcane, we identified nine H2B histones as expressed in internodes (Table 3). Sequence analysis revealed high similarity among these histones, with differences concentrated in the N-terminal tail, a characteristic common in plant nucleosomes (Chabouté *et al*., 1993). Little is known about H2B variants, with only the isoforms H2BFWT and H2B1A expressed in human testes, which have been studied in depth to date (Molden *et al*., 2015). This study also identified several PTMs in H2B, including H2B.6K32acR33me1 and H2B.K3me3 (Figure 3). In animals, H2BK32 is part of the N-terminal HBR (H2B repression domain), with lysines critical for transcriptional repression by stabilising the interaction between H2B and DNA (Parra *et al*., 2006). However, given the dissimilarity of the N-terminal tail between animals and plants, it is unclear whether a functional parallel exists, mainly because the HBR domain corresponds to the region near lysine 50 in sugarcane H2B. H2B.K3me3 has been previously described in *Neurospora crassa* (Anderson *et al*., 2010).

For histone H4, PTMs were identified across the entire protein, with special emphasis on the central domain (Figure 4). These PTMs add new insights to those previously reported for sugarcane (Moraes *et al*., 2015). Until now, PTMs identified in sugarcane H4 were limited to the N-terminal tail. This difference can be attributed to the higher sensitivity of the methods employed in this study and the more significant number of LC-MS/MS runs performed. The modifications found in the central and C-terminal domains of H4 have been reported in animals (Luense *et al*., 2016; Wang *et al*., 2019) but remain understudied in plants. Although several PTMs in H4 were identified, our study was unable to quantify these modifications efficiently in sugarcane H4. This limitation may be related to the experimental design, which was adapted from a protocol optimised for H3. Improvements in chromatography and sample preparation are likely necessary for a more detailed study of H4 PTMs.

The PTMs identified in H3 were similar to those previously described in the literature (Figure 5). Moraes et al. (2015) reported that H3 appears to have acetyl groups limited to the N-terminal tail. However, studies using the ChIP technique have demonstrated that H3K56ac, located in the central region of the histone, is present in *Arabidopsis* (Roudier *et al*., 2011). Thus, the method used in this study appears insensitive to this modification, a conclusion supported by other studies that also failed to detect acetylation in the central domain of H3 using mass spectrometry-based techniques (Zhang *et al*., 2007; Wu *et al*., 2009). In our study, the most abundant and easily detectable modifications in H3 were methylation and acetylation at lysines 9, 14, 17, 18, 23, 26, 27, 36, and 37. Regarding mono- and dimethylation in arginines, we detected modifications at residues 8, 17, 26, and 40, with the latter PTM being exclusively identified in H3.3d (Figure 1). Among the observed changes in H3, the peptide corresponding to residues 27–40 contains modifications that differ between H3.1 and H3.3 (Table 2). This extensive diversity of modifications likely reflects the importance of this region for the functions of these proteins.

In sorghum, H3K4me3 and H3K36me3 were correlated with active transcription sites, while H3K27me2/me3 and H3K36me2 correlated with inactive sites (Zhou *et al*., 2021). We did not identify H3K4me3, only me1, in this study (Figure 5); however, this modification has been identified in previous studies (Moraes *et al*., 2015). The most abundant PTMs in H3.1 were K27me1/2 (Figure 6c and Table 2). Methylation of lysine 27 on histone H3.1 (H3.1K27me) by the Polycomb Repressive Complex 2 (PRC2) is a hallmark of facultative heterochromatin in various organisms (Wiles & Selker, 2017). In plants, PRC1 and especially PRC2 operate interdependently depending on the genomic context to ensure transcriptional repression in heterochromatic regions (Wang & Shen, 2018).). K27me3 was also frequently found (Figure 6c and Table 2). In *Saccharum spontaneum*, the H3K27 demethylase activity of the JmjC protein from the aKDM4/JHDM3 group represses the drought response (Yu *et al*., 2024). Thus, our results for sugarcane align with observations in other organisms, as the most abundant PTMs in H3.1 are repressive marks. However, our study revealed lower levels of H3.1K36me1 than previously reported (Moraes *et al*., 2015), while the modification H3.1K27ac was detected at higher levels(Figure 6c), which could be the result of the growth conditions in each experiment (i., controlled versus field conditions).

H3.3 has distinct functions from H3.1 in *Arabidopsis* (Jacob *et al*., 2014). While H3.3 is associated with actively expressed genes in animals and plants, H3.1 is incorporated into the genome during replication and is associated with transcriptionally inactive regions (Hake & Allis, 2006; Jacob *et al..*, 2014; Jiang & Berger, 2017). Chromatin immunoprecipitation (ChIP) studies have demonstrated that H3.3 associates with gene bodies, the regions between transcription start and end sites, and exhibits a positive correlation with transcription. It has been proposed that H3.3 inhibits chromatin compaction mediated by H1, allowing methyltransferases access to DNA and leading to gene body methylation (Wollmann *et al*., 2017).

Among the PTMs exclusively identified for H3.3 (peptide 27–40), we found that K36me2K37me1 was the most abundant PTM, representing approximately 30% of the 24 PTM combinations identified (Figure 6d). Following this, K36me1/2/3 and K36ac each accounted for around 10%. While H3K36 PTMs are conserved active gene markers across various organisms, plants utilise biological mechanisms distinct from those in animals. It has been demonstrated that K36me2/3 marks active transcription of genes involved in flowering time regulation and other processes in *Arabidopsis* (Xu *et al*., 2008). The primary function of K36me2 is linked to transcriptional elongation, and in plants, this modification plays a similar role to that of K36me3 in animals. Additionally, H3K36me2 is highly correlated with m6A modifications and mRNA in plants, underscoring its importance in co-regulating mRNA modifications and chromatin context during transcription (Shim *et al*., 2020).

H3K36me3 is associated with transcriptional elongation in animals and yeast, peaking at the 3’ end of genes. Conversely, H3K36ac in yeast is primarily found at promoters. In plants, H3K36me3 peaks at the 5’ half of genes, while H3K36ac is confined to approximately 480 bp distal from the transcription start site. In *Arabidopsis*, H3K36ac and H3K36me3 overlap downstream of the transcription start site, with H3K36ac showing levels inversely correlated with H3K36me3 (Mahrez *et al*., 2016). Our study revealed that in sugarcane, a significant portion of K36ac co-occurs with K37me1, whereas K36me3 is predominantly found alone (Figure 6c and d). These findings demonstrate that PTMs in H3.1 and H3.3 are largely evolutionarily conserved.

We also identified mono- and dimethylation at H3.3R40 (Figure 5). These PTMs were described in human cells (Shi *et al*., 2017), and their functions remain largely unexplored. Considering that the structural difference between H3.1 and H3.3 consists of only four amino acids, we hypothesise that H3.3R40me1/2 may play roles specific to the functions of this histone. Other vital modifications include H3.1K9 and H3.1K14 in the peptide corresponding to residues 9–26 in H3.1a (Figure 5). H3K9me3 is present in heterochromatin in metazoans, whereas in plants, H3K9me2 fulfills this role (Feng & Jacobsen, 2011).

This study identified and quantified methylation and acetylation at H3.1K9 and H3.1K14 (Figure 6a). The most abundant modifications observed were H3K14ac and H3K9me1, primarily found in isolation, likely due to their antagonistic functions (Figure 6a). Our findings are similar to those previously reported for sugarcane, except for H3K9me2, where our study demonstrated a lower relative abundance (Moraes *et al*., 2015). This difference supports the hypothesis that the abundance of H3K9me2 may vary depending on experimental conditions.

## Conclusions

This work represents the first comprehensive analysis of histone PTMs for H2A and H2B in sugarcane. We have mapped a detailed profile of these modifications, identified heavily modified sites and quantified their relative abundances. Our research expands the current knowledge about sugarcane’s epigenetic landscape, which could aid in understanding of gene regulation, drive improvement strategies, and enhance stress adaptation approaches in this crop.

## Acknowledgements

This work received financial support from the São Paulo Research Foundation (FAPESP) under grant numbers 11/00818-8 and 22/13970-7 as part of the BIOEN Program. JPLS received a Conselho Nacional de Desenvolvimento Científico e Tecnológico (CNPq) scholarship. We thank Professors Paolo Di Mascio and Graziella E. Ronsein for the Mass spectrometry analyses performed at the Redox Proteomics Core of the Mass Spectrometry Resource at Chemistry Institute, University of Sao Paulo (FAPESP grant numbers 2012/12663-1, 2023/00995-4, CEPID Redoxoma 2013/07937-8), and Dr Mariana P. Massafera for her technical assistance.

## Author Contributions

Conceptualisation, JPLS, JACM, and CTH; Methodology, JPLS, JACM, ARMS, MSC, JC; Software, JPLS, and CTH; Formal Analysis, JPLS; Investigation, JPLS, FMAJ, ARMS and CTH; Resources, MSC and CTH; Data Curation, JPLS; Writing – Original Draft Preparation, JPLS and CTH; Writing – Review & Editing, JPLS, ACM, FMAJ and CTH; Visualization, CTH; Supervision, CTH; Project Administration, CTH; Funding Acquisition, CTH.

## Conflict of Interest and Other Ethics Statements

The authors declare no conflicts.

## References

Anderson DC, Green GR, Smith K, Selker EU. 2010. Extensive and Varied Modifications in Histone H2B of Wild-Type and Histone Deacetylase 1 Mutant Neurospora crassa. Biochemistry 49: 5244–5257.

van Attikum H, Gasser SM. 2009. Crosstalk between histone modifications during the DNA damage response. Trends in Cell Biology 19: 207–217.

Berr A, Shafiq S, Shen W-H. 2011. Histone modifications in transcriptional activation during plant development. Biochimica et Biophysica Acta (BBA) - Gene Regulatory Mechanisms 1809: 567–576.

Bowler C, Benvenuto G, Laflamme P, Molino D, Probst AV, Tariq M, Paszkowski J. 2004. Chromatin techniques for plant cells. The Plant Journal: For Cell and Molecular Biology 39: 776–789.

Chabouté ME, Chaubet N, Gigot C, Philipps G. 1993. Histones and histone genes in higher plants: Structure and genomic organization. Biochimie 75: 523–531.

Cheavegatti-Gianotto A, de Abreu HMC, Arruda P, Bespalhok Filho JC, Burnquist WL, Creste S, di Ciero L, Ferro JA, de Oliveira Figueira AV, de Sousa Filgueiras T, et al. 2011. Sugarcane (Saccharum X officinarum): A Reference Study for the Regulation of Genetically Modified Cultivars in Brazil. Tropical Plant Biology 4: 62–89.

Chen D, Yu G, Wu X, Ye M, Wang Q. 2024. H3K4 demethylase SsJMJ11 promotes flowering in sugarcane. Industrial Crops and Products 216: 118718.

Dal-Bianco M, Carneiro MS, Hotta CT, Chapola RG, Hoffmann HP, Garcia AA, Souza GM. 2012. Sugarcane improvement: how far can we go? Curr Opin Biotechnol 23: 265–70.

D’Hont A, Grivet L, Feldmann P, Glaszmann JC, Rao S, Berding N. 1996. Characterisation of the double genome structure of modern sugarcane cultivars (Saccharum spp.) by molecular cytogenetics. Molecular and General Genetics MGG 250: 405–413.

Earley KW, Shook MS, Brower-Toland B, Hicks L, Pikaard CS. 2007. In vitro specificities of Arabidopsis co-activator histone acetyltransferases: implications for histone hyperacetylation in gene activation. The Plant Journal 52: 615–626.

Feng S, Jacobsen SE. 2011. Epigenetic modifications in plants: an evolutionary perspective. Current Opinion in Plant Biology 14: 179–186.

Garcia BA, Mollah S, Ueberheide BM, Busby SA, Muratore TL, Shabanowitz J, Hunt DF. 2007. Chemical derivatization of histones for facilitated analysis by mass spectrometry. Nature protocols 2: 933–938.

Garsmeur O, Droc G, Antonise R, Grimwood J, Potier B, Aitken K, Jenkins J, Martin G, Charron C, Hervouet C, et al. 2018. A mosaic monoploid reference sequence for the highly complex genome of sugarcane. Nature Communications 9: 2638.

Giaimo BD, Ferrante F, Herchenröther A, Hake SB, Borggrefe T. 2019. The histone variant H2A.Z in gene regulation. Epigenetics & Chromatin 12: 37.

Gómez-Zambrano Á, Merini W, Calonje M. 2019. The repressive role of Arabidopsis H2A.Z in transcriptional regulation depends on AtBMI1 activity. Nature Communications 10: 2828.

Hake SB, Allis CD. 2006. Histone H3 variants and their potential role in indexing mammalian genomes: The “H3 barcode hypothesis”. Proceedings of the National Academy of Sciences 103: 6428–6435.

Healey AL, Garsmeur O, Lovell JT, Shengquiang S, Sreedasyam A, Jenkins J, Plott CB, Piperidis N, Pompidor N, Llaca V, et al. 2024. The complex polyploid genome architecture of sugarcane. Nature 628: 804–810.

Henikoff S, Smith MM. 2015. Histone Variants and Epigenetics. Cold Spring Harbor Perspectives in Biology 7: a019364.

Horn V, Uckelmann M, Zhang H, Eerland J, Aarsman I, le Paige UB, Davidovich C, Sixma TK, van Ingen H. 2019. Structural basis of specific H2A K13/K15 ubiquitination by RNF168. Nature Communications 10: 1751.

Hu Y, Chen X, Zhou C, He Z, Shen X. 2022. Genome-wide identification of chromatin regulators in Sorghum bicolor. 3 Biotech 12: 117.

Jacob Y, Bergamin E, Donoghue MTA, Mongeon V, LeBlanc C, Voigt P, Underwood CJ, Brunzelle JS, Michaels SD, Reinberg D, et al. 2014. Selective Methylation of Histone H3 Variant H3.1 Regulates Heterochromatin Replication. Science 343: 1249–1253.

Jiang D, Berger F. 2017. Histone variants in plant transcriptional regulation. Biochimica et Biophysica Acta (BBA) - Gene Regulatory Mechanisms 1860: 123–130.

Joseph FM, Young NL. 2023. Histone variant-specific post-translational modifications. Seminars in Cell & Developmental Biology 135: 73–84.

Kawashima T, Lorković ZJ, Nishihama R, Ishizaki K, Axelsson E, Yelagandula R, Kohchi T, Berger F. 2015. Diversification of histone H2A variants during plant evolution. Trends in Plant Science 20: 419–425.

Kouzarides T. 2007. Chromatin Modifications and Their Function. Cell 128: 693–705.

Kumar SV. 2018. H2A.Z at the Core of Transcriptional Regulation in Plants. Molecular Plant 11: 1112–1114.

Luense LJ, Wang X, Schon SB, Weller AH, Lin Shiao E, Bryant JM, Bartolomei MS, Coutifaris C, Garcia BA, Berger SL. 2016. Comprehensive analysis of histone post-translational modifications in mouse and human male germ cells. Epigenetics & Chromatin 9: 24.

MacLean B, Tomazela DM, Shulman N, Chambers M, Finney GL, Frewen B, Kern R, Tabb DL, Liebler DC, MacCoss MJ. 2010. Skyline: an open source document editor for creating and analyzing targeted proteomics experiments. Bioinformatics 26: 966–968.

Mahrez W, Arellano MST, Moreno-Romero J, Nakamura M, Shu H, Nanni P, Köhler C, Gruissem W, Hennig L. 2016. H3K36ac Is an Evolutionary Conserved Plant Histone Modification That Marks Active Genes. Plant Physiology 170: 1566– 1577.

McCormick RF, Truong SK, Sreedasyam A, Jenkins J, Shu S, Sims D, Kennedy M, Amirebrahimi M, Weers BD, McKinley B, et al. 2018. The Sorghum bicolor reference genome: improved assembly, gene annotations, a transcriptome atlas, and signatures of genome organization. The Plant Journal: For Cell and Molecular Biology 93: 338–354.

Molden RC, Bhanu NV, LeRoy G, Arnaudo AM, Garcia BA. 2015. Multi-faceted quantitative proteomics analysis of histone H2B isoforms and their modifications. Epigenetics & Chromatin 8: 15.

Moraes I, Yuan Z-F, Liu S, Souza GM, Garcia BA, Casas-Mollano JA. 2015. Analysis of Histones H3 and H4 Reveals Novel and Conserved Post-Translational Modifications in Sugarcane. PloS One 10: e0134586.

Nunez-Vazquez R, Desvoyes B, Gutierrez C. 2022. Histone variants and modifications during abiotic stress response. Frontiers in Plant Science 13.

Osakabe A, Lorković ZJ, Kobayashi W, Tachiwana H, Yelagandula R, Kurumizaka H, Berger F. 2018. Histone H2A variants confer specific properties to nucleosomes and impact on chromatin accessibility. Nucleic Acids Research 46: 7675–7685.

Parra MA, Kerr D, Fahy D, Pouchnik DJ, Wyrick JJ. 2006. Deciphering the Roles of the Histone H2B N-Terminal Domain in Genome-Wide Transcription. Molecular and Cellular Biology 26: 3842–3852.

Piperidis N, D’Hont A. 2020. Sugarcane genome architecture decrypted with chromosome-specific oligo probes. The Plant Journal 103: 2039–2051.

Roudier F, Ahmed I, Bérard C, Sarazin A, Mary-Huard T, Cortijo S, Bouyer D, Caillieux E, Duvernois-Berthet E, Al-Shikhley L, et al. 2011. Integrative epigenomic mapping defines four main chromatin states in Arabidopsis. The EMBO journal 30: 1928–1938.

Shi L, Wen H, Shi X. 2017. The histone variant H3.3 in transcriptional regulation and human disease. Journal of molecular biology 429: 1934–1945.

Shim S, Lee HG, Lee H, Seo PJ. 2020. H3K36me2 is highly correlated with m6A modifications in plants. Journal of Integrative Plant Biology 62: 1455–1460.

Strahl BD, Allis CD. 2000. The language of covalent histone modifications. Nature 403: 41–45.

Sura W, Kabza M, Karlowski WM, Bieluszewski T, Kus-Slowinska M, Pawełoszek Ł, Sadowski J, Ziolkowski PA. 2017. Dual Role of the Histone Variant H2A.Z in Transcriptional Regulation of Stress-Response Genes. The Plant Cell 29: 791–807.

Tafrova JI, Tafrov ST. 2014. Human histone acetyltransferase 1 (Hat1) acetylates lysine 5 of histone H2A in vivo. Molecular and Cellular Biochemistry 392: 259–272.

Tyanova S, Temu T, Cox J. 2016. The MaxQuant computational platform for mass spectrometry-based shotgun proteomics. Nature Protocols 11: 2301–2319.

Vettore AL, Silva FR da, Kemper EL, Souza GM, Silva AM da, Ferro MIT, Henrique-Silva F, Giglioti ÉA, Lemos MVF, Coutinho LL, et al. 2003. Analysis and Functional Annotation of an Expressed Sequence Tag Collection for Tropical Crop Sugarcane. Genome Research 13: 2725–2735.

Vivek Hari Sundar G, Madhu A, Archana A, Shivaprasad PV. 2024. Plant histone variants at the nexus of chromatin readouts, stress and development. Biochimica et Biophysica Acta (BBA) -General Subjects 1868: 130539.

Wang T, Gao H, Li W, Liu C. 2019. Essential Role of Histone Replacement and Modifications in Male Fertility. Frontiers in Genetics 10.

Wang M, Li A-M, Chen Z-L, Qin C-X, Liao F, Pan Y-Q, Lakshmanan P, Li X-F, Huang D-L. 2025. Dynamic proteome and acetylome profiling reveals key regulators of sucrose accumulation in sugarcane. Plant Cell Reports 44: 74.

Wang Q, Shen W-H. 2018. Chromatin modulation and gene regulation in plants: insight about PRC1 function. Biochemical Society Transactions 46: 957–966.

Wiles ET, Selker EU. 2017. H3K27 methylation: a promiscuous repressive chromatin mark. Current Opinion in Genetics & Development 43: 31–37.

Wollmann H, Stroud H, Yelagandula R, Tarutani Y, Jiang D, Jing L, Jamge B, Takeuchi H, Holec S, Nie X, et al. 2017. The histone H3 variant H3.3 regulates gene body DNA methylation in Arabidopsis thaliana. Genome Biology 18: 94.

Wu T, Yuan T, Tsai S-N, Wang C, Sun S-M, Lam H-M, Ngai S-M. 2009. Mass spectrometry analysis of the variants of histone H3 and H4 of soybean and their post-translational modifications. BMC Plant Biology 9: 98.

Xiong L, Wang Y. 2011. Mapping post-translational modifications of histones H2A, H2B and H4 in Schizosaccharomyces pombe. International Journal of Mass Spectrometry 301: 159–165.

Xu L, Zhao Z, Dong A, Soubigou-Taconnat L, Renou J-P, Steinmetz A, Shen W-H. 2008. Di- and Tri- but Not Monomethylation on Histone H3 Lysine 36 Marks Active Transcription of Genes Involved in Flowering Time Regulation and Other Processes in Arabidopsis thaliana. Molecular and Cellular Biology 28: 1348–1360.

Yelagandula R, Stroud H, Holec S, Zhou K, Feng S, Zhong X, Muthurajan UM, Nie X, Kawashima T, Groth M, et al. 2014. The Histone Variant H2A.W Defines Heterochromatin and Promotes Chromatin Condensation in *Arabidopsis*. Cell 158: 98– 109.

Yu G, Chen D, Ye M, Wu X, Zhu Z, Shen Y, Mehareb EM, Esh A, Raza G, Wang K, et al. 2024. H3K27 demethylase SsJMJ4 negatively regulates drought-stress responses in sugarcane. Journal of Experimental Botany 75: 3040–3053.

Zhang K, Sridhar VV, Zhu J, Kapoor A, Zhu J-K. 2007. Distinctive Core Histone Post-Translational Modification Patterns in Arabidopsis thaliana. PLOS ONE 2: e1210.

Zhou C, Zhou H, Ma X, Yang H, Wang P, Wang G, Zheng L, Zhang Y, Liu X. 2021. Genome-Wide Identification and Characterization of Main Histone Modifications in Sorghum Decipher Regulatory Mechanisms Involved by mRNA and Long Noncoding RNA Genes. Journal of Agricultural and Food Chemistry 69: 2337–2347.

