## Supplemental Table for "Mass Spectrometry-Based Characterization of Histone Post-Translational Modifications in Sugarcane"

**Supplemental Table 1 – Chromosomal distribution of sorghum and sugarcane histones.** Sorghum and sugarcane chromosomes were paired following Healey et al. (2024). The histone with the highest frequency in each chromosome is in bold.

|  |  | **H1** | **H2A** | **H2B** | **CENH3** | **H31** | **H33** | **H33L** | **H4** |
| --- | --- | --- | --- | --- | --- | --- | --- | --- | --- |
| **Chr. A** | **sorghum 1** | 1 | **7** | 1 | 0 | 0 | 1 | 2 | 2 |
|  | **sugarcane 1** | 7 | **31** | 6 | 0 | 1 | 5 | 13 | 10 |
| **Chr. B** | **sorghum 2** | 1 | **4** | 3 | 0 | 0 | 0 | 1 | 2 |
|  | **sugarcane 5** | 7 | **31** | 23 | 0 | 0 | 0 | 4 | 23 |
| **Chr. C** | **sorghum 3** | 0 | 0 | **5** | 0 | 2 | 0 | 0 | 4 |
|  | **sugarcane 2** | 0 | 0 | **25** | 0 | 11 | 0 | 0 | 8 |
| **Chr. D** | **sorghum 4** | 0 | 1 | 1 | 0 | 1 | 1 | 0 | 1 |
|  | **sugarcane 3** | 0 | 6 | 6 | 0 | **8** | 4 | 0 | 4 |
| **Chr. E** | **sorghum 5** | 0 | **2** | 0 | 0 | 0 | 0 | 0 | 0 |
|  | **sugarcane 10** | 0 | **5** | 0 | 0 | 0 | 0 | 0 | 1 |
| **Chr. F** | **sorghum 6** | 1 | 0 | 0 | 0 | **2** | 1 | 0 | 1 |
|  | **sugarcane 7** | 5 | 0 | 0 | 0 | 8 | **9** | 0 | 5 |
| **Chr. G** | **sorghum 7** | 0 | 0 | **2** | 0 | 0 | 0 | 0 | 0 |
|  | **sugarcane 8** | 0 | 1 | **8** | 0 | 0 | 0 | 0 | 0 |
| **Chr. H** | **sorghum 8** | 0 | **2** | 0 | 0 | 0 | 0 | 0 | 0 |
|  | **sugarcane 9** | 0 | **5** | 0 | 0 | 0 | 0 | 0 | 1 |
| **Chr. I** | **sorghum 9** | 0 | **2** | 1 | 1 | 1 | 0 | 0 | 1 |
|  | **sugarcane 6** | 0 | **12** | 0 | 5 | 0 | 0 | 0 | 4 |
| **Chr. J** | **sorghum 10** | 1 | 0 | 0 | 0 | **2** | 1 | 0 | 0 |
|  | **sugarcane 4** | 4 | 0 | 0 | 0 | **8** | 5 | 1 | 0 |

**Supplemental Table 2 – Catalog of histones H1 genes found in sorghum and sugarcane.** Sorghum bicolor genome v5.1, Sugarcane R570 genome, and the SUCEST Database (Sugarcane Assembled Sequences, SAS).

| Histone | Sorghum locus | Sugarcane locus | SAS |
| --- | --- | --- | --- |
| H1.1 | Sobic.006G186700 | SoffiXsponR570.07Ag170300  SoffiXsponR570.07Bg182300  SoffiXsponR570.07Cg180800  SoffiXsponR570.07Dg170700  SoffiXsponR570.07Eg131000  SoffiXsponR570.7_10Ag286700 | SCBFRZ2018H11.g  SCCCRZ2C04H07.g |
| H1.2 | Sobic.002G056300 | SoffiXsponR570.01Ag292000  SoffiXsponR570.01Bg275500  SoffiXsponR570.01Cg142000  SoffiXsponR570.05Ag067300  SoffiXsponR570.05Bg057800  SoffiXsponR570.05Cg057400  SoffiXsponR570.05Dg059300  SoffiXsponR570.05Eg052200  SoffiXsponR570.05Fg031800  SoffiXsponR570.05Gg060700  SoffiXsponR570.1Z292200  SoffiXsponR570.5_9Ag366900 | SCCCLB1001E09.g  SCCCLR2004E01.g  SCEPRZ1008C04.g  SCRUFL1113E08.g  SCRUFL1120G08.g |
| H1.3 | Sobic.010G021300 | SoffiXsponR570.04Ag032000  SoffiXsponR570.04Bg023300  SoffiXsponR570.04Dg026700  SoffiXsponR570.04Gg023600 | SCACLR2014A04.g  SCAGFL3022A04.g  SCBGLR1119A06.g  SCCCLR2C02H09.g  SCQSLR1018A10.g |
| H1.3 | Sobic.001G054800 | SoffiXsponR570.01Ag055100  SoffiXsponR570.01Bg052300  SoffiXsponR570.01Cg027200  SoffiXsponR570.01Eg056100 | SCACFL8019G10.g  SCBGLR1082F03.g  SCCCLR1079F01.g  SCCCRZ2002H02.g  SCJFRZ2010B03.g  SCRUFL1015E09.g  SCRUFL1020A01.g |

**Supplemental Table 3 – Catalog of histones H2A genes found in sorghum and sugarcane.** *Sorghum bicolor* genome v5.1, Sugarcane R570 genome, and the SUCEST Database (Sugarcane Assembled Sequences, SAS).

| Histone | Sorghum locus | Sugarcane locus | SAS |
| --- | --- | --- | --- |
| H2A1 | Sobic.002G330000 | SoffiXsponR570.05Ag293900  SoffiXsponR570.05Bg284600  SoffiXsponR570.05Cg289400  SoffiXsponR570.05Dg294200  SoffiXsponR570.05Eg286100  SoffiXsponR570.05Fg220300  SoffiXsponR570.5_9Ag105400 | SCJFRZ2010D04.g  SCJFRZ2014A08.g  SCVPLR2019A02.g |
| H2A2 | Sobic.002G329900 | SoffiXsponR570.05Ag293700  SoffiXsponR570.05Bg284500  SoffiXsponR570.05Dg294100  SoffiXsponR570.05Eg286000  SoffiXsponR570.05Fg220200  SoffiXsponR570.5_9Ag105500 | SCJLLR2013B11.g  SCJLLR1101F06.g |
| H2A.X1 | Sobic.001G110000 | SoffiXsponR570.01Ag114800  SoffiXsponR570.01Bg105100  SoffiXsponR570.01Eg114500  SoffiXsponR570.01Eg114900 | SCBFST3135H08.g |
| H2A.X2 | Sobic.008G114500 | SoffiXsponR570.09Ag101700  SoffiXsponR570.09Cg101400  SoffiXsponR570.09Dg048700  SoffiXsponR570.09Dg055900  SoffiXsponR570.5_9Ag467200  SoffiXsponR570.9us90g014900 | SCCCLR1068C05.g |
| H2A.X3 | Sobic.001G296700 | SoffiXsponR570.01Ag300300  SoffiXsponR570.01Bg277900  SoffiXsponR570.01Cg147600  SoffiXsponR570.01Dg159000  SoffiXsponR570.01Fg099000 | SCCCLR1001G05.g  SCEPAM1020H08.g |
| H2A.W1  H2A.W2  H2A.W5 | Sobic.001G009700  Sobic.002G059200  Sobic.002G276000 | SoffiXsponR570.01Ag009700  SoffiXsponR570.01Bg010200  SoffiXsponR570.01Eg009600  SoffiXsponR570.05Ag070900  SoffiXsponR570.05Ag242300  SoffiXsponR570.05Bg061000  SoffiXsponR570.05Bg229800  SoffiXsponR570.05Cg060600  SoffiXsponR570.05Cg235600  SoffiXsponR570.05Dg061900  SoffiXsponR570.05Dg248100  SoffiXsponR570.05Eg055700  SoffiXsponR570.05Eg056100  SoffiXsponR570.05Eg232400  SoffiXsponR570.05Fg032800  SoffiXsponR570.05Fg169200  SoffiXsponR570.05Gg064100  SoffiXsponR570.5_9Ag155100  SoffiXsponR570.5_9Ag363600  SoffiXsponR570.8_5Ag000100 | SCCCLR2001G07.g  SCAGLR1021A12.g  SCCCLR1001C03.g  SCCCRZ2C03C04.g  SCUTLR2008G03.g |
| H2A.W3  H2A.W4 | Sobic.001G416800  Sobic.001G416900 | SoffiXsponR570.01Ag419200  SoffiXsponR570.01Ag419500  SoffiXsponR570.01Bg375700  SoffiXsponR570.01Bg375800  SoffiXsponR570.01Cg266900  SoffiXsponR570.01Cg267000  SoffiXsponR570.01Dg273700  SoffiXsponR570.01Dg273800  SoffiXsponR570.01Eg327400  SoffiXsponR570.01Eg327700  SoffiXsponR570.01Fg214900  SoffiXsponR570.01Fg215000 | SCCCLR1048F07.g  SCCCFL3003C11.g  SCCCLB1024F05.g  SCCCLR2C02C12.g  SCEZAM1082G06.g |
| H2A.W6 | Sobic.009G015900 | SoffiXsponR570.06Ag011200  SoffiXsponR570.06Eg018600  SoffiXsponR570.06Fg015400  SoffiXsponR570.6_9Ag107600  SoffiXsponR570.6us88g048100 | SCCCLR2001C09.g |
| H2A.W7 | Sobic.009G164900 | SoffiXsponR570.06Ag134700  SoffiXsponR570.06Cg111200  SoffiXsponR570.06Dg083800  SoffiXsponR570.06Eg143100  SoffiXsponR570.06Fg082300  SoffiXsponR570.06Gg066500  SoffiXsponR570.6_9Ag245900 | SCCCRZ2002H03.g |
| H2A.W8 | Sobic.008G126100 |  |  |
| H2A.Z1 | Sobic.004G205300 | SoffiXsponR570.01Eg404000  SoffiXsponR570.03Ag188500  SoffiXsponR570.03Bg170100  SoffiXsponR570.03Cg181400  SoffiXsponR570.03Dg197000  SoffiXsponR570.03Eg155200  SoffiXsponR570.03Eg155300 | SCCCCL4007C09.g |
| H2A.Z2 | Sobic.001G094500 | SoffiXsponR570.01Ag098400  SoffiXsponR570.01Bg090700  SoffiXsponR570.01Dg022600  SoffiXsponR570.01Eg098700  SoffiXsponR570.05Dg342800 | SCEPLR1008D05.g  SCJLLR1101A02.g |
| H2A.Z3 | Sobic.005G168500 | SoffiXsponR5700Ag156300  SoffiXsponR5700Bg155100  SoffiXsponR5700Cg141900  SoffiXsponR5700Eg075300  SoffiXsponR5700Fg048100 |  |
| H2A.Z4 | Sobic.001G495700 | SoffiXsponR570.01Bg451700  SoffiXsponR570.01Eg403900 |  |
| H2A.ZL1 | Sobic.005G082900 |  |  |
| H2A.ZL2 | - |  |  |

**Supplemental Table 4 – Catalog of histones H2B genes found in sorghum and sugarcane.** *Sorghum bicolor* genome v5.1, Sugarcane R570 genome, and the SUCEST Database (Sugarcane Assembled Sequences, SAS).

| Histone | Sorghum locus | Sugarcane locus | SAS |
| --- | --- | --- | --- |
| H2B1  H2B2  H2B3  H2B4  H2B5  H2B7  H2B8  H2B11  H2B10  H2B12  H2B13 | Sobic.003G200501  Sobic.002G210200  Sobic.004G265400  Sobic.003G067300  Sobic.002G403400  Sobic.003G067500  Sobic.009G164200  Sobic.003G092200  Sobic.007G221800  Sobic.007G149600  Sobic.002G138000 | SoffiXsponR570.02Ag068200  SoffiXsponR570.02Ag068300  SoffiXsponR570.02Ag185900  SoffiXsponR570.02Bg065300  SoffiXsponR570.02Bg065500  SoffiXsponR570.02Bg154600  SoffiXsponR570.02Cg066900  SoffiXsponR570.02Cg179100  SoffiXsponR570.02Dg065100  SoffiXsponR570.02Dg065200  SoffiXsponR570.02Dg183100  SoffiXsponR570.02Eg044400  SoffiXsponR570.02Eg152200  SoffiXsponR570.02Eg152300  SoffiXsponR570.02Fg066800  SoffiXsponR570.02Fg067100  SoffiXsponR570.02Fg166200  SoffiXsponR570.02Gg084700  SoffiXsponR570.03Ag257100  SoffiXsponR570.03Bg228400  SoffiXsponR570.03Cg286800  SoffiXsponR570.03Dg259800  SoffiXsponR570.03Eg262400  SoffiXsponR570.03Gg121900  SoffiXsponR570.05Ag134400  SoffiXsponR570.05Ag179800  SoffiXsponR570.05Ag365400  SoffiXsponR570.05Bg120300  SoffiXsponR570.05Bg169500  SoffiXsponR570.05Bg362000  SoffiXsponR570.05Cg124200  SoffiXsponR570.05Cg174900  SoffiXsponR570.05Cg175300  SoffiXsponR570.05Cg363000  SoffiXsponR570.05Dg185800  SoffiXsponR570.05Dg363800  SoffiXsponR570.05Eg108300  SoffiXsponR570.05Eg166700  SoffiXsponR570.05Eg336700  SoffiXsponR570.05Fg050200  SoffiXsponR570.05Fg104300  SoffiXsponR570.05Fg297500  SoffiXsponR570.05Gg128900  SoffiXsponR570.05Gg191500  SoffiXsponR570.08Ag215500  SoffiXsponR570.08Bg140400  SoffiXsponR570.08Cg127200  SoffiXsponR570.08Dg169500  SoffiXsponR570.1Z005000  SoffiXsponR570.1Z223400  SoffiXsponR570.5_9Ag031700  SoffiXsponR570.5_9Ag217300  SoffiXsponR570.5_9Ag295800  SoffiXsponR570.8_10Ag264500  SoffiXsponR570.8_10Ag339900  SoffiXsponR570.8_5Ag142900  SoffiXsponR570.8_5Ag244800 | SCBGFL3095E08.g  SCBGLR1002E03.g  SCCCLR1024A09.g  SCCCLR1048C02.g  SCCCLR1068G11.g  SCCCLR1C04B01.g  SCCCLR2C02B06.g  SCCCRZ1001H04.g  SCEPLB1042A02.g  SCJLLR1101H08.g  SCJLLR2020F01.g  SCMCRT2108A05.g  SCQGLR1041C10.g  SCQGLR1062G09.g  SCSFFL3090D03.g  SCSGAM1096G11.g  SCUTLR2015D03.g |
| H2B9 | Sobic.003G350100 | SoffiXsponR570.02Ag335900  SoffiXsponR570.02Bg300200  SoffiXsponR570.02Bg300800  SoffiXsponR570.02Cg327300  SoffiXsponR570.02Dg326400  SoffiXsponR570.02Eg300200  SoffiXsponR570.02Gg231200 | SCCCLR2002G11.g |
| H2B6 | Sobic.001G417000 | SoffiXsponR570.01Ag419600  SoffiXsponR570.01Bg375900  SoffiXsponR570.01Cg267100  SoffiXsponR570.01Dg273900  SoffiXsponR570.01Eg327800  SoffiXsponR570.01Fg215100 | SCEZLB1014E02.g  SCCCRZ1004F04.g |

**Supplemental Table 5 – Catalog of histones CENH3 genes found in sorghum and sugarcane.** *Sorghum bicolor* genome v5.1, Sugarcane R570 genome, and the SUCEST Database (Sugarcane Assembled Sequences, SAS).

| Histone | Sorghum locus | Sugarcane locus | SAS |
| --- | --- | --- | --- |
| CENH3 | Sobic.009G178900 | SoffiXsponR570.06Ag148600  SoffiXsponR570.06Cg124300  SoffiXsponR570.06Dg099200  SoffiXsponR570.06Gg081100  SoffiXsponR570.6_9Ag259800 | SCCCLR2004A05.g |

**Supplemental Table 6 – Catalog of histones H3.1 genes found in sorghum and sugarcane.** *Sorghum bicolor* genome v5.1, Sugarcane R570 genome, and the SUCEST Database (Sugarcane Assembled Sequences, SAS).

| Histone | Sorghum locus | Sugarcane locus | SAS |
| --- | --- | --- | --- |
| H3.1a  H3.1c  H3.1e  H3.1b  H3.1g  H3.1h | Sobic.010G047100  Sobic.006G081800  Sobic.010G047200  Sobic.004G170500  Sobic.009G152902  Sobic.003G367402 | SoffiXsponR570.02Bg317900  SoffiXsponR570.02Cg344400  SoffiXsponR570.02Dg304800  SoffiXsponR570.02Eg317900  SoffiXsponR570.02Fg313600  SoffiXsponR570.02Gg247700  SoffiXsponR570.03Ag155800  SoffiXsponR570.03Ag156000  SoffiXsponR570.03Bg136900  SoffiXsponR570.03Cg151100  SoffiXsponR570.03Cg151200  SoffiXsponR570.03Dg163900  SoffiXsponR570.03Eg123200  SoffiXsponR570.03Gg039000  SoffiXsponR570.04Ag059200  SoffiXsponR570.04Ag059300  SoffiXsponR570.04Bg050500  SoffiXsponR570.04Cg052800  SoffiXsponR570.04Cg052900  SoffiXsponR570.04Gg052900  SoffiXsponR570.4us91g047600  SoffiXsponR570.4us91g047900 | SCCCLR1001F12.g  SCCCLR1068F12.g  SCCCLR1C04B07.g  SCCCLR2001D10.g  SCCCRZ2001G08.g  SCEQLB1066C12.g  SCMCRT2102E07.g  SCQGLR1019F06.g  SCVPLB1016B08.g  SCVPLB1017H05.g |
| H3.1d | Sobic.006G076400 | SoffiXsponR570.01Eg314700  SoffiXsponR570.07Ag067300  SoffiXsponR570.07Ag067800  SoffiXsponR570.07Bg075100  SoffiXsponR570.07Cg071900  SoffiXsponR570.07Dg062100  SoffiXsponR570.07Dg065600  SoffiXsponR570.7_10Ag177900  SoffiXsponR570.7_10Ag178000 | SCAGLR2026E02.g  SCBFLR1026A06.g  SCBGLR1002H11.g  SCEQLB1063F02.g |
| H3.1f | Sobic.003G065002 | SoffiXsponR570.02Dg063800  SoffiXsponR570.02Eg043000  SoffiXsponR570.02Fg065200  SoffiXsponR570.02Cg064700  SoffiXsponR570.02Ag066500 | SCAGFL8013E01.g  SCJFRZ2009C12.g  SCMCLR1032D12.g  SCQGLR1062C03.g |

**Supplemental Table 7 – Catalog of histones H3.3 genes found in sorghum and sugarcane.** *Sorghum bicolor* genome v5.1, Sugarcane R570 genome, and the SUCEST Database (Sugarcane Assembled Sequences, SAS).

| Histone | Sorghum locus | Sugarcane locus | SAS |
| --- | --- | --- | --- |
| H3.3a | Sobic.006G100600 | SoffiXsponR570.07Ag087800  SoffiXsponR570.07Bg096300  SoffiXsponR570.07Cg096000  SoffiXsponR570.07Cg096500  SoffiXsponR570.07Dg112100  SoffiXsponR570.07Eg043100  SoffiXsponR570.7_10Ag200300 | SCCCLR2001F08.g |
| H3.3b | Sobic.010G021402 | SoffiXsponR570.04Ag032100  SoffiXsponR570.04Bg023400  SoffiXsponR570.04Cg024700  SoffiXsponR570.04Dg026800  SoffiXsponR570.04Gg023700  SoffiXsponR570.07Cg213800  SoffiXsponR570.7_10Ag319100 | SCBGLR1002A05.g  SCBGLR1023F03.g  SCCCCL7C02A12.g  SCCCFL5003D06.g  SCJFLR1035H03.g  SCJFRT1060F02.g  SCJFRZ2006G04.g  SCRFLR1055C06.g  SCUTLR2008H04.g  SCVPRZ2039D03.g |
| H3.3c | Sobic.001G350501 | SoffiXsponR570.01Ag352200  SoffiXsponR570.01Bg316200  SoffiXsponR570.01Dg209900  SoffiXsponR570.01Eg265400  SoffiXsponR570.01Fg150800 | SCJFLR1013C11.g  SCSGFL4037A02.g |
| H3.3d | Sobic.004G188000 | SoffiXsponR570.03Ag171500  SoffiXsponR570.03Dg178800  SoffiXsponR570.03Eg139200  SoffiXsponR570.03Gg053700 | SCBGFL3095F09.g  SCCCCL3120E10.g  SCEQLR1094C02.g |

**Supplemental Table 8 – Catalog of histones H3-like genes found in sorghum and sugarcane.** *Sorghum bicolor* genome v5.1, Sugarcane R570 genome, and the SUCEST Database (Sugarcane Assembled Sequences, SAS).

| Histone | Sorghum locus | Sugarcane locus | SAS |
| --- | --- | --- | --- |
| H3L1 | Sobic.001G297400 | SoffiXsponR570.01Ag300800  SoffiXsponR570.01Cg148100  SoffiXsponR570.01Dg159500  SoffiXsponR570.01Eg214900  SoffiXsponR570.01Fg099500 | - |
| H3L2  H3L3 | Sobic.001G284800  Sobic.002G355700 | SoffiXsponR570.01Ag287400  SoffiXsponR570.01Bg271200  SoffiXsponR570.01Cg137900  SoffiXsponR570.01Dg147900  SoffiXsponR570.01Dg148000  SoffiXsponR570.01Dg148300  SoffiXsponR570.01Fg087200  SoffiXsponR570.05Ag320100  SoffiXsponR570.05Cg315300  SoffiXsponR570.05Fg247600  SoffiXsponR570.1Z287700  SoffiXsponR570.4us91g063500  SoffiXsponR570.5_9Ag079900 | - |

**Supplemental Table 9 – Catalog of histones H4 genes found in sorghum and sugarcane.** *Sorghum bicolor* genome v5.1, Sugarcane R570 genome, and the SUCEST Database (Sugarcane Assembled Sequences, SAS).

| Histone | Sorghum locus | Sugarcane locus | SAS |
| --- | --- | --- | --- |
| H4.1 | Sobic.002G291066 | SoffiXsponR570.05Ag256100  SoffiXsponR570.05Bg244700  SoffiXsponR570.05Cg250400  SoffiXsponR570.05Dg260100  SoffiXsponR570.05Eg246200  SoffiXsponR570.05Fg182200  SoffiXsponR570.5_9Ag142300 | SCCCRZ1004C05.g  SCJLHR1028C12.g  SCSBFL1043F08.g  SCSGHR1071E06.g |
| H4.2 | Sobic.003G057367 | SoffiXsponR570.02Ag060300  SoffiXsponR570.02Bg057700  SoffiXsponR570.02Cg058700  SoffiXsponR570.02Cg058700  SoffiXsponR570.02Dg057500  SoffiXsponR570.02Fg058800  SoffiXsponR570.02Fg058900 | SCEPLR1051H12.g  SCJFLR1013A12.g  SCJLLR2020B06.g |
| H4.3 | Sobic.003G056600 | - | - |
| H4.4 | Sobic.003G347900 | SoffiXsponR570.02Bg297800  SoffiXsponR570.02Gg228700 | SCCCRZ2001G01.g  SCJLLR1011E11.g |
| H4.5 | Sobic.006G191800 | SoffiXsponR570.07Ag175000  SoffiXsponR570.07Cg185200  SoffiXsponR570.07Dg175900  SoffiXsponR570.07Eg135700 | SCCCLR2002G09.g |
| H4.6 | Sobic.001G313200 | SoffiXsponR570.01Ag315200  SoffiXsponR570.01Bg294400  SoffiXsponR570.01Cg163900  SoffiXsponR570.01Dg176300  SoffiXsponR570.01Eg229500  SoffiXsponR570.01Fg114900  SoffiXsponR570.10Ag081600.1  SoffiXsponR570.7_10Ag147100  SoffiXsponR570.9us90g015500 | SCCCLR2001D12.g  SCJFRZ2009F02.g |
| H4.7 | Sobic.002G210500 | SoffiXsponR570.05Ag180100  SoffiXsponR570.05Ag180300  SoffiXsponR570.05Bg169800  SoffiXsponR570.05Bg170000  SoffiXsponR570.05Cg175200  SoffiXsponR570.05Cg175600  SoffiXsponR570.05Cg175900  SoffiXsponR570.05Dg186200  SoffiXsponR570.05Eg167000  SoffiXsponR570.05Eg167200  SoffiXsponR570.05Fg104700  SoffiXsponR570.05Fg104900  SoffiXsponR570.05Gg191800  SoffiXsponR570.05Gg192200  SoffiXsponR570.05_9Ag216800  SoffiXsponR570.05_9Ag217000 | SCCCLR2003E01.g  SCSBLB1033H09.g |
| H4.8 | Sobic.001G529400 | SoffiXsponR570.01Bg484900  SoffiXsponR570.01Cg381300  SoffiXsponR570.01Dg381500  SoffiXsponR570.01Fg326100 | SCAGLB1070C07.g  SCAGLR2026F05.g  SCSGRT2062A10.g  SCUTLR2015B06.g |
| H4.9 | Sobic.003G057100 | - | SCCCLR1078H02.g  SCRFLR2038C05.g |
| H4.10 | Sobic.009G167500 | SoffiXsponR570.06Dg086700  SoffiXsponR570.06Eg146000  SoffiXsponR570.06Fg085000  SoffiXsponR570.06Gg069600 | SCCCLR2002D04.g |
| H4.11 | Sobic.004G279601 | SoffiXsponR570.03Ag233500  SoffiXsponR570.03Cg230500  SoffiXsponR570.03Dg245700  SoffiXsponR570.03Eg275500 | SCCCLR2001D01.g  SCVPRZ2035C06.g |
